# Enabling high-plex spectral imaging via DNA-barcoded signal tuning and panel optimization

**DOI:** 10.64898/2026.03.18.709053

**Authors:** Robert Reinhardt, Tatjana Straka, Wouter-Michiel Vierdag, Kristina Jevdokimenko, Frank Hecht, Elena Pianfetti, Tim Hudelmaier, Hoyin Lai, Wernher Fouquet, Florian Fahrbach, M. Julia Roberti, Anna Kreshuk, Sinem K. Saka

## Abstract

High-plex spectral imaging has the potential to transform the analysis of spatial organization in cells and tissues, yet its practical implementation remains limited by challenges in panel design, sample preparation, signal balancing, and experimental validation. While cyclic imaging approaches are widely used in spatial omics, spectral imaging across the full fluorescence spectrum and computational unmixing remain underutilized due to these challenges. Here, we present a generalizable framework for high-plex spectral imaging that leverages DNA-barcoded labeling and programmable signal amplification to provide precise control over fluorescence signal composition. Orthogonal DNA barcodes decouple target labeling from fluorophore detection, enabling reversible fluorophore application and systematic panel optimization directly on the same sample. Programmable DNA-based amplification further enables independent and quantitative tuning of fluorescence intensities across targets, overcoming a key limitation of spectral unmixing, namely imbalanced signal contributions in overlapping channels, and thereby improving accuracy and robustness. The framework also supports the generation of experiment-specific ground truth datasets and systematic evaluation of unmixing algorithms, providing a quantitative basis for panel validation and performance assessment. We demonstrate the practical implementation of this framework by developing a panel for simultaneous imaging of 15 subcellular structures without fluidic cycling and using the optimized panel to profile the effects of chemical perturbations on subcellular organization. We quantitatively evaluate panel compilation and provide a rigorous assessment unmixing performance using both linear and reference-free unmixing methods. Importantly, we leverage foundation models trained on standard fluorescence data, for segmentation-free, high-dimensional analysis of spectrally unmixed images without needing large datasets or model retraining. Together, we establish a practical and tunable framework for high-plex spectral imaging that lowers experimental barriers and enables broader adoption of spectral unmixing for biological and biomedical applications.

## Main Text

The rise of single-cell sequencing and growing sets of analysis tools for high-dimensional data has created great interest in increasing the number of features that can be detected from the same cells. The field of spatial omics has been booming to meet this demand by enabling extraction of a high number of features without losing the *in situ* context of cells and tissues. For fluorescence imaging-based spatial omics methods, one of the most critical enabling factors was the development of approaches for cyclic detection which allowed overcoming the spectral overlap limitation by reuse of the same spectrally non-overlapping fluorescent channels in sequential cycles of labeling, imaging, and erasing of the signal^1^. Although there are different implementations, cyclic labeling or detection is typically a common requirement for highly multiplexed fluorescence imaging. Well-established or commercialized imaging-based spatial omics methods typically (with few exceptions, such as^2^) rely only on the well-separated 3-5 spectral channels that are re-utilized through tens of fluidic cycling rounds.

On the other hand, multispectral imaging with spectral unmixing approaches have been around for a long time and have a very high potential to boost multiplexing levels without cycles of relabeling and imaging^3–6^. Yet the full range of the fluorescence spectrum have been rarely exploited for high-plex *in situ* barcoding and detection, largely owing to major practical limitations in implementing spectral unmixing reliably on biological samples. For example, typically, spectral unmixing necessitates the use of narrower wavelength bands to minimize the overlap between different spectral channels. This significantly decreases the signal-to-noise ratio, and in the presence of noise the unmixing-related errors become more pronounced as the spectral overlap of the fluorophores increases^7^.

Achieving reliable high-plex unmixing experiments require optimizations on multiple fronts. On the instrument side, device components, and acquisitions settings need to be set right to minimize noise and crosstalk for the given panel of fluorophores, including the use of low noise detectors and photon counting mode, sequential imaging of channels with optimal arrangement of lasers and detection windows^8^. On the panel design side optimizing the fluorophore-target matches become critical to avoid unbalanced signal distribution and extensive spatial overlap of targets for spectrally overlapping fluorophores. And on the image processing side, it is important to choose a suitable spectral unmixing algorithm and validate the performance to obtain reliable results. These requirements have made mainstream adoption of unmixing for fluorescence imaging challenging. Although a small number of recent studies achieved dedicated ≥10-plex panels^9–11^, the complexity of these experiments tends to increase greatly with every additional channel.

Recent improvements in imaging technology and practical hardware and software solutions simplify setting up spectral imaging experiments^12–18^, and new analysis paradigms such as growing number of algorithms for reference-free, also known as ”blind” unmixing^11,17,19–24^ reduce the complexity of data acquisition (e.g., the need to have single-plex reference images for each fluorophore to perform linear unmixing) with more scalable data processing. However, benchmarking and panel generation standards, ground truths (GT), references, and evaluation metrics^25^ are still critically missing, and unmixing performance for fluorophore panels beyond 10-plex have not been characterized in detail. Most studies rely on synthetic validation data that does not account for sample-related challenges, such as signal and target distribution complexities, sample heterogeneity and endogenous fluorescence (either intrinsic or arising from sample fixation/embedding). Spectral imaging also typically requires high signal for reliable separation and detection efficiency, hence would benefit significantly from incorporation of tunable in situ signal amplification methods. To address these constraints, here we repurpose a previously established open source method for DNA-barcoding and fluidic exchange-based multiplexed imaging and signal amplification, namely SABER^26,27^.

Immuno-SABER is a DNA-based signal amplification strategy for highly multiplexed protein imaging in which antibodies are conjugated to DNA barcodes which are used as docks for single-stranded DNA concatemers generated by primer exchange reaction (PER)^28^. PER is an isothermal DNA polymerization mechanism in which a primer is repeatedly extended on a hairpin template, generating programmable single-stranded DNA concatemers without enzymatic cycling or thermal denaturation. PER enables control over sequence, length, and amplification gain. PER-generated concatemers (typically extended to 500-600 nucleotides) act as amplifiers by offering repetitive binding sites for short fluorophore-conjugated oligonucleotides (“imagers”). Iterative hybridization and dehybridization of imagers to DNA concatemers enable repeated, multiplexed readout of many targets using a limited set of fluorophores. By decoupling amplification from fluorescence readout, Immuno-SABER supports spatially resolved detection of many proteins in cells and tissues using standard fluorescence microscopy and cyclic imaging workflows. If desired, concatemers can also be used as branching sites for secondary concatemers to amplify the signal further.

Even though it was originally designed for cyclic imaging, SABER offers unique advantages for high-plex multispectral imaging. Here, we are using two capabilities offered by SABER to develop a generalizable validation platform for high-plex spectral unmixing panels: (i) the cyclic DNA-exchange capability to optimize the fluorophore-to-target matching and to generate controls, unmixing references, and GT images for assessment of the performance of the panel and unmixing approaches directly on the same cells, (ii) independent tunability of signal amplification to increase the signal-to-noise and to balance the contribution of crosstalking fluorophores when needed under image acquisition conditions that maximize the unmixing performance. Using these features, biological targets and fluorophore panels can be set almost independently as the target labeling and fluorescence detection steps are uncoupled via orthogonal DNA barcoding of the primary probes. We aimed to leverage this flexibility for developing a generalizable panel building workflow for establishing and validating bespoke high-plex panels.

We demonstrate this workflow on a custom panel targeting main subcellular structures (primarily membranous and non-membranous organelle markers) that can be used as a visual profiling panel for subcellular organization. Using this panel, we generate high-resolution 3D imaging data with a point scanning confocal microscope and quantitatively characterize the unmixing performance of linear and reference-free unmixing directly for our panel and sample. We show that tunable signal amplification is instrumental to increase signal to noise and signal to crosstalk for problematic fluorophores and critically improves the overall performance of the panel. Finally, we utilize the panel and spectral unmixing to perform 15-plex subcellular profiling for two perturbations without relying on lengthy fluidic exchange steps. Applying a recent foundation model, which was trained on traditional 3- or 4-plex images, we can detect the altered spatial organization of the targets in response to these perturbations in our high-plex spectrally unmixed images. Hence, we establish the feasibility of using our spectral imaging workflow for high-plex and high-resolution imaging, and provide data that can be further used as a reference for multiplexed organelle labeling. Our results open the path for future high-throughput subcellular profiling experiments that leverage spectral unmixing for high-plex data generation.

## Results

For developing our workflow, we focused on a panel targeting main subcellular structures (primarily membranous and non-membranous organelle markers) that can be used as a visual profiling panel for subcellular organization. The panel covers a diverse selection of subcellular distributions (spatially overlapping, non-overlapping, spot-like, filamentous, and diffuse), intensities, and microenvironments (membrane, cytoplasm, nuclear, nucleolar, mitochondrial, condensate etc.) and hence provides example cases for different challenges regarding spectral unmixing (**Fig. 1a**).

**Figure 1.**
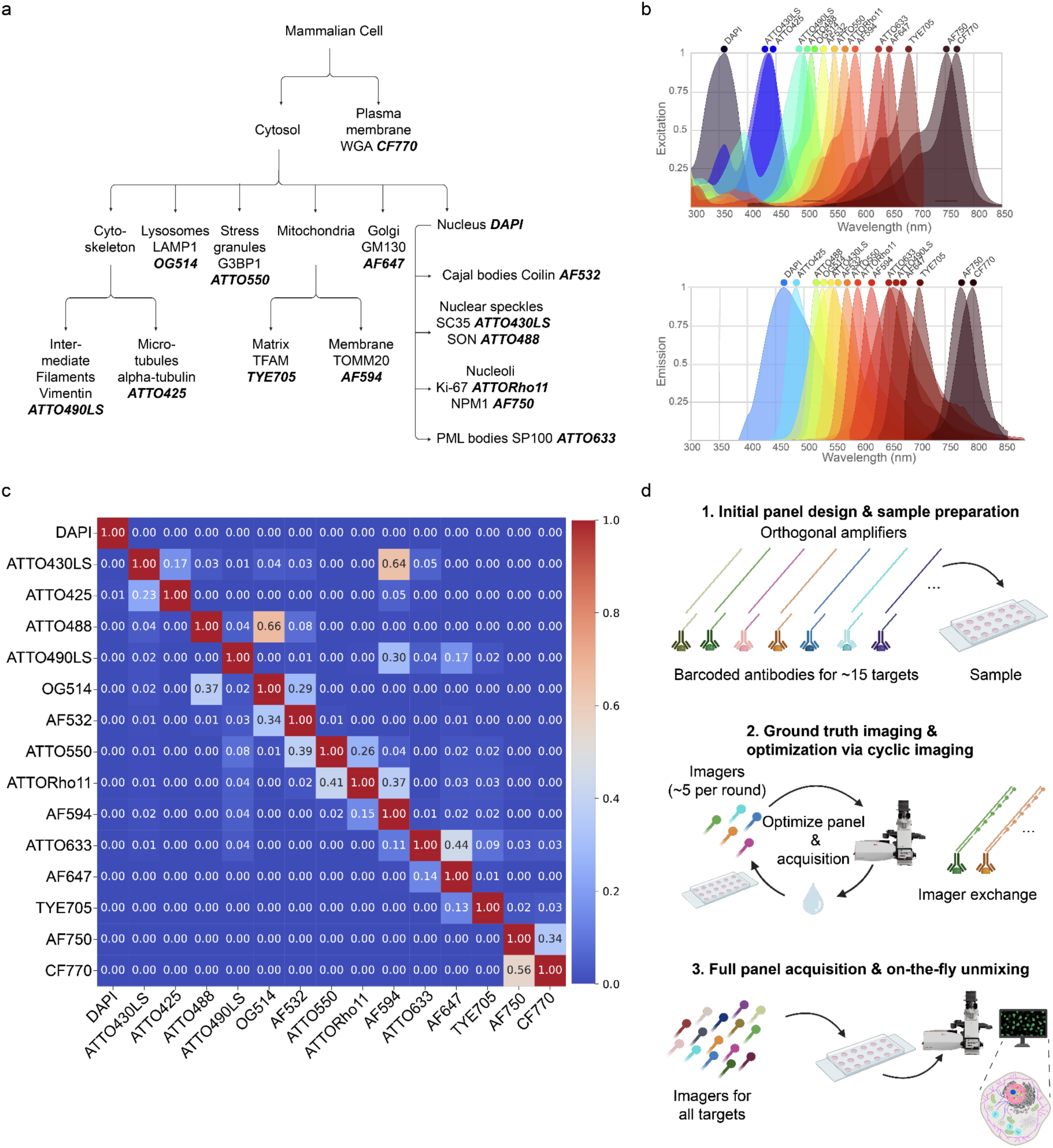
Overview of the organellar targets and dye panel. **(a)** Subcellular panel and assigned fluorescent dye. **(b)** Excitation and emission spectra of selected dyes (obtained from fpbase.org^37,38^). **(c)** Theoretical crosstalk of selected dye panel (obtained from LAS X 3D viewer software). **(d)** Summary of ground truth acquisition with iterative group approach. The process starts with an optimized Immuno-SABER protocol using a full antibody and concatemer panel. Subsequent cycles employ spectrally distinct dyes for validation, followed by the single round multiplexed imaging of all targets to obtain the raw data for spectral unmixing.

To create a large spectral multiplexing panel for a one-shot high-plex approach, we have tested >20 potential fluorophores. In addition to fluorophores commonly used for immunofluorescence (such as ATTO488 or AF647), we have tested a set of non-conventional fluorophores that are spectrally well-distributed to make use of the visible and near-infrared spectrum (400-800 nm) (**Supplementary File 1 - Tab 1**). After the initial in situ characterization, we settled on 15 main fluorophores (referred to later as 15-plex, **Fig. 1b-c**) that showed favorable characteristics for a high-plex panel such as high photostability, low unspecific labeling, and high brightness in aqueous imaging media (**Supplementary File 1 - Tab 1** and **Supplementary File 2 - Tab 1**).

For assigning the fluorophores to the targets in the panel, it is advisable to consider the expected spatial distribution pattern and expression level of the targets, such that the most spectrally overlapping fluorophores are assigned to least spatially overlapping targets (**Fig. 1a-c**), and weak fluorophores are matched to higher abundance targets as much as possible. It is important to note that sufficient information for fluorophore-to-target matching may not always be available a priori when building a new panel, underscoring the need to have a validation platform for building such panels.

For initial establishment of the panel, we utilize cyclic imaging to apply smaller subgroups of spectrally distinct fluorophores iteratively on the same sample. This approach allows us to validate the panel and generate GT data before performing the full multiplex experiment in a single round of imaging (**Fig. 1d**). SABER relies on fluorescently tagged DNA probes (imagers) that are hybridized on demand to the barcoded amplifier sequences on the targets, allowing efficient decoupling of labeling from imaging readout. This permits quick re-shuffling of the target-fluorophore combinations. After characterizing the panel with these initial imaging rounds, we dehybridize the last set of imagers and proceed to perform the multiplexed imaging round where all imagers are present at the same time.

First, we assigned orthogonal oligo barcodes to marker proteins that are known to primarily localize to the target structures^29^ and also included DAPI stain for nuclear labeling (see **Supplementary Note 1** regarding use of DAPI in spectral imaging panels), and fluorophore-coupled lectin, wheat-germ-agglutinin (WGA), as a membrane-directed stain. To increase the ease of antibody-DNA barcoding and to decrease unspecific background of DNA-barcoded antibodies and oligonucleotide probes, we made multiple improvements to the previous Immuno-SABER protocol^27^: (i) utilization of adaptor-based site-specific conjugation of antibodies instead of non-targeted conjugation of lysines, (ii) increased oligo blocking and temporary double-stranding of barcodes, (iii) blocking of any potential crosstalk between imagers in high-plex panels by use of decoy imagers (**Supplementary Fig. S1**). As further detailed in the **Methods** section and partially in recent work^30^, these changes improve the robustness not only for Immuno-SABER but are also applicable for any other method that utilizes DNA-barcoded antibodies and oligo probes (for e.g.^9,31–36)^.

As a starting point of the panel validation framework, using the reference spectra and theoretical crosstalk predictions (**Fig. 1b-c),** we have divided our fluorophores into 3 sets of 4-5 spectrally distinct channels plus DAPI, which is included in each round and was used for alignment of images across cycles (**Fig. 2a-c**). This way, we can generate the GT images for each target by performing only 2 exchange rounds (**Fig. 2d**). The GT data is not only instrumental for validation but is also utilized to assess the performance of the assigned target-fluorophore matches and generate benchmarks for signal intensity and spatial distribution of the targets, so that the panel and signal intensities can be further finetuned for the best performance. Furthermore, GT images can be also used as references for linear unmixing algorithms (as demonstrated before^14^).

**Figure 2.**
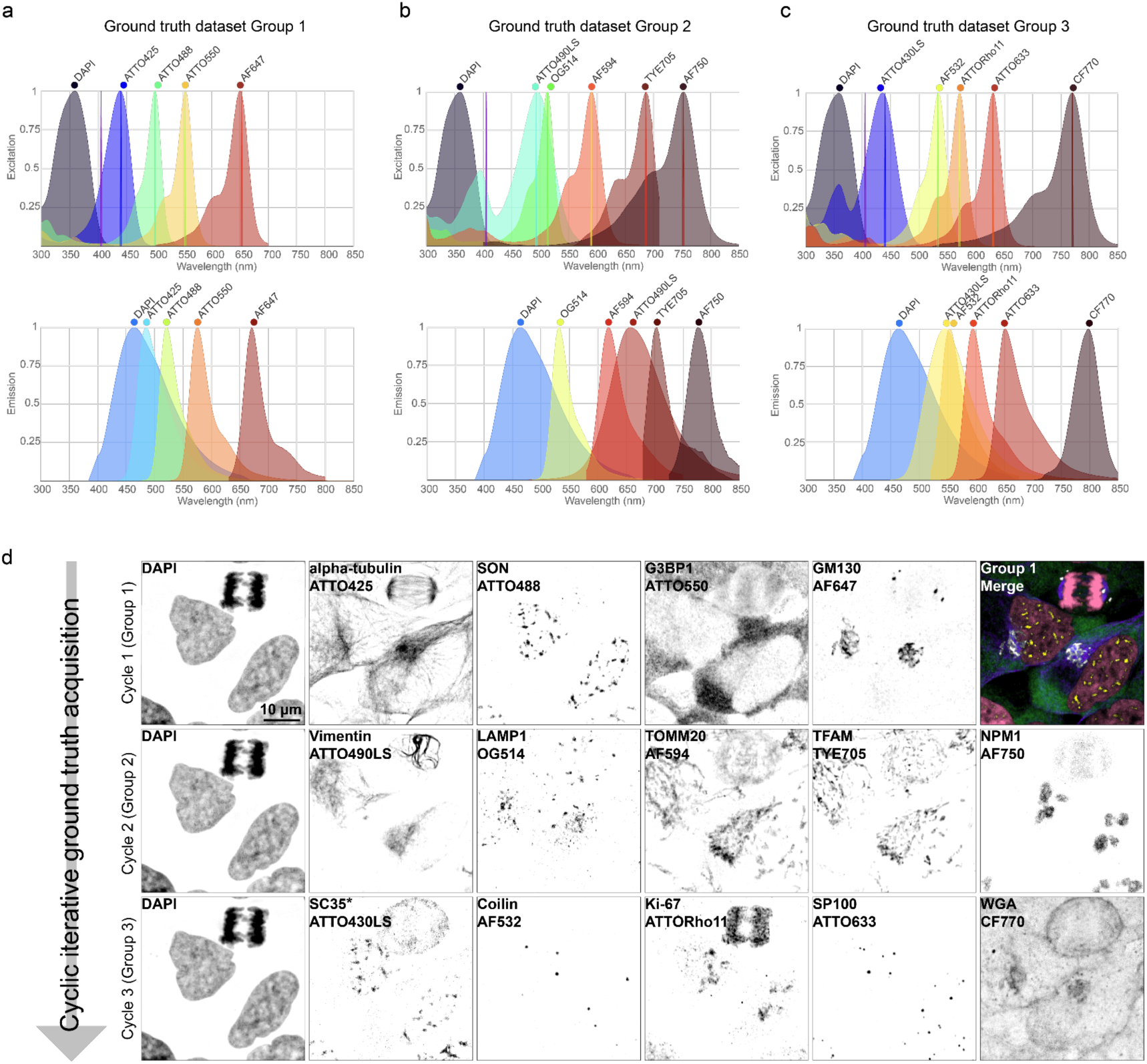
Group approach and ground truth (GT) validation. **(a-c)** The entire panel is separated into individual groups of spectrally non-overlapping dyes. Excitation and emission spectra were obtained from fpbase.org^37,38^. **(d)** Results of three Immuno-SABER cycles with spectrally distinct fluorescent dyes, imaged with optimal settings for 4- to 5-plex imaging (extended depth of field projection of 3D confocal stacks). An example overlay image is shown for the Group 1 dyes. SC35-ATTO430LS(*) underwent branched Immuno-SABER to compensate for the weak signal yielded by this antibody-fluorophore pair. Acquisition settings are described in **Supplementary File 2 - Tab 2**.

For this optimization round, signal for each target was amplified by use of ssDNA concatemers that are 500-600 nucleotides (nt) long. One of the fluorophores, the long Stokes shift dye ATTO430LS yielded relatively low signal, particularly when imaged under settings that are optimized for 15-plex spectral unmixing. Hence, we concluded that this fluorophore should ideally be used in conjunction with more abundant targets or with higher signal amplification (for example by branching of SABER concatemers for exponential signal amplification, where secondary concatemers of 300-350 nt are applied on top of the primary concatemers). Hence, after the initial cyclic imaging round of groups (**Fig. 2d**), we removed the ATTO430LS from the multiplex panel and moved forward with the other 14 markers.

The acquired 14-plex spectrally mixed images (“raw”) were then subjected to unmixing (**Fig. 3a**). We applied either linear unmixing or a reference-free unmixing algorithm that would be considered semi-blind, as it does not use reference images but incorporates input spectra along with system and setup information. To evaluate the effect of unmixing on the GT dataset, we considered two main points: how the representation of the real signal improves after unmixing (self-to-self comparison), and how the spectral crosstalk between different channels decreases. We considered two commonly used metrics to evaluate these two points: structural similarity index measure (SSIM)^39^ and Pearson’s correlation coefficient (PCC). After obtaining SSIM and PCC values for GT vs. GT comparison a few differences between the metrics became apparent. SSIM, one of the most popular metrics for analyzing spectrally unmixed images, was useful to evaluate the self-to-self similarity, however it was uninformative to evaluate the spectral crosstalk between different channels (**Supplementary Fig. S2**). Because SSIM quantification considers features such as luminance, contrast and structure between two images, it becomes difficult to interpret SSIM values as a performance measure. We noticed that different cellular markers might return high similarity values although they represent very different organelles. For example, comparing Ki-67 (nuclear marker) and GM130 (Golgi apparatus marker) we observed high SSIM (SSIM = 0.7), which does not reflect their distribution profile and low biological co-localization. SSIM’s sensitivity was also previously shown to be affected by signal to noise ratio alterations^40^.

**Figure 3.**
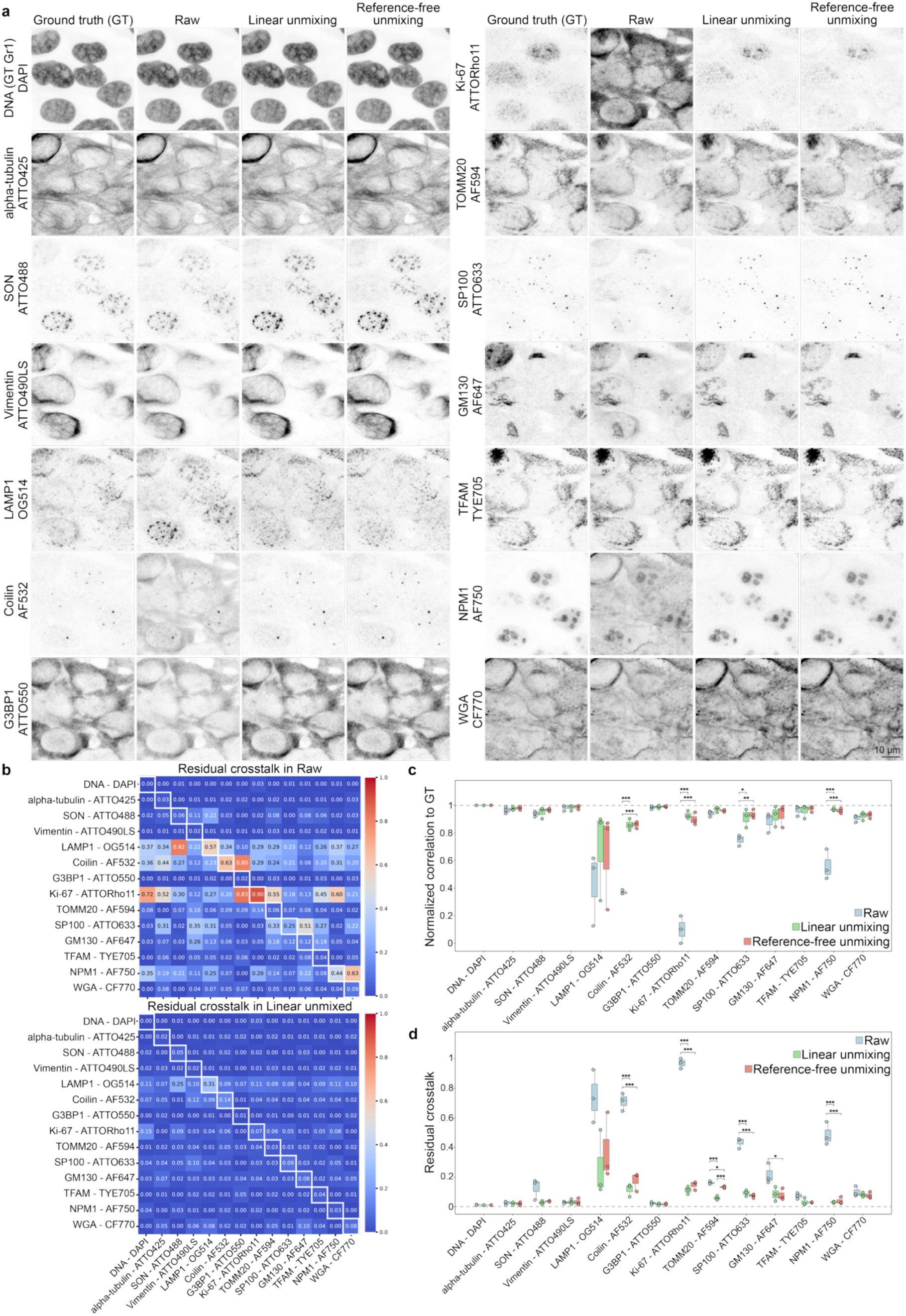
Quantitative assessment of spectral unmixing performance. **(a)** Representative images from the GT, raw (spectrally mixed), linear unmixed, and reference-free unmixed 14-plex datasets. **(b)** Residual crosstalk heatmaps calculated from raw and linear unmixed datasets accounted for endogenous/natural correlation of different markers in GT dataset. For each channel comparison mean values from 3 fields of views (FOVs) are shown. **(c-d)** The box-and-whisker plot with individual data points displays data from n = 3 FOVs. The center line represents the mean, boxes indicate the 25th and 75th percentiles, and whiskers extend to the minimum and maximum values. Significance was measured using a one-way ANOVA with Tukey’s HSD post-hoc test (p < 0.05 = *, p < 0.01 = **, p < 0.001 = ***). In panel **c**, normalized correlation to GT increases to over 0.8 and in panel **d** total residual crosstalk decreases to below 0.2 after both linear and reference-free unmixing, with the exception of LAMP1-OG514, where the improvement provided by unmixing is more modest. Acquisition settings are described in **Supplementary File 2 - Tab 2** and **3**.

In contrast, PCC measures the linear correlation between the pixel intensity patterns and hence is expected to be more informative to evaluate preservation of subcellular distributions. Using PCC, we first observed naturally occurring biological correlations in GT vs. GT (**Supplementary Fig. S3, Supplementary Fig. S4a**). For example, two mitochondrial markers TFAM and TOMM20 showed high PCC as expected (PCC = 0.87). Further examples include nuclear markers such as DAPI and Ki-67 (PCC = 0.72) and Ki-67 and NPM1 (PCC = 0.60). At the same time, nucleus- and cytosol-residing proteins showed negative correlation. For example, DAPI and G3BP1 showed a strong negative correlation (PCC = -0.44), NPM1 and G3BP1 as well (PCC = -0.3), and Ki-67 and G3BP1 (PCC = -0.28). Ki-67 and GM130, previously showing high SSIM (**Supplementary Fig. S2**), resulted in low correlation (PCC = 0.07) as expected. Overall, PCC values accurately reflect the expected localization overlap between the different structures, and hence PCC is a more aligned to intuitive human evaluation and is a more suitable metric than SSIM to quantify the unmixing performance for cell biology, where target signal localization is a critical feature of the images.

We then proceeded with evaluating the self-to-self PCC improvement after unmixing, which showed how similar the same images were to the GT dataset after unmixing. To do this, we compared same channel PCC values for raw vs. GT, linear-unmixed vs. GT and reference-free unmixed vs. GT datasets (**Fig. 3c, Supplementary Fig. S4b**). For the majority of the channels normalized PCC was already high as expected based on the low theoretical crosstalk with other channels (**Fig. 1c**) and thus improvement after linear and reference-free unmixing was minimal (DNA-DAPI, α-tubulin-ATTO425, SON-ATTO488, Vimentin-ATTO490LS, G3BP1-ATTO550, TOMM20-AF594, GM130-AF647, TFAM-TYE705, WGA-CF770).

There were, however, 5 channels where we observed high crosstalk and low correlation to GT (PCC<0.8 down to 0.1). For 4 of these channels, both linear and reference-free unmixing significantly improved the correlation with GT (Coilin-AF532, Ki-67-ATTORho11, SP100-ATTO633, NPM1-AF750). The most problematic channel was LAMP1-OG514 which showed low self-to-self correlation and high residual crosstalk after linear and reference-free unmixing. We aimed to identify which channels contributed most to the crosstalk. Having the GT vs. GT dataset comparisons, we were able to normalize our observed correlations for naturally occurring biological colocalization between each marker pair. To do this, we obtained PCC values for raw vs. GT, linear-unmixed vs. GT and reference-free unmixed vs. GT datasets and subtracted GT vs. GT comparisons from these matrices to achieve what we refer to as “residual spectral crosstalk” (**Fig. 3b**, **Supplementary Fig. S4c**). Upon this detailed inspection we were able to conclude that in the unmixed images the correct LAMP1-OG514 signal was being masked by surrounding channels (particularly SON-ATTO488, can also be visualized in the images in **Fig. 3a**). To summarize the impact of crosstalk for each channel in a more simplified manner, we added and normalized all the residual crosstalk values and plotted the total residual crosstalk (**Fig. 3d**). Here we could see that the residual crosstalk mimics the trend with self-to-self correlation, confirming that spectral crosstalk was robustly accounted for and the fluorophores were correctly unmixed. Overall, linear unmixing showed less residual crosstalk than reference-free unmixing method, albeit this difference being statistically significant only for one marker (TOMM20-AF594).

### Panel optimization with tunable signal amplification

The use of PCC with the GT images and residual crosstalk which accounts for natural colocalization in the panel are instrumental to evaluate the panel and unmixing performance and identify problematic channels. Based on these results, we sought to further optimize our multiplex panel. LAMP1 signal was rather low in the GT images due to OG514 fluorophore being in a spectrally dense region with other fluorophores and having one of the narrowest detection windows (17 nm) in the panel. We hypothesized that tuning the signal intensity for this channel by branched Immuno-SABER amplification would improve the accuracy of unmixing (**Fig. 4a**). When Immuno-SABER with single-concatemer amplification (×1) was applied, LAMP1-OG514 signal in the raw images was almost half of the crosstalk from SON-ATTO488 into this channel (**Fig. 4b-c**), yielding a low ratio of signal to crosstalk both in raw and resulting unmixed images. This quantification reflects the low linear unmixing performance revealed by our residual crosstalk analysis (SON-ATTO488 to LAMP1-OG514 had the highest residual crosstalk as shown in **Fig. 3b**). Branched SABER amplification (×2) evened out the SON-ATTO488 and LAMP1-OG514 signals in raw images and substantially improved (∼4-fold) the signal to crosstalk ratio for LAMP1-OG514 both in raw and unmixed images. On the opposite side, amplification of the LAMP1 signal naturally decreased the originally very high SON-ATTO488 signal to the crosstalk from neighboring LAMP1-OG514. However, the ATTO488 signal was still double the crosstalk from OG514 in raw images and yielded satisfactory unmixing results (**Fig. 4b-c**), hence the final panel was optimized to include branched amplification for LAMP1-OG514.

**Figure 4.**
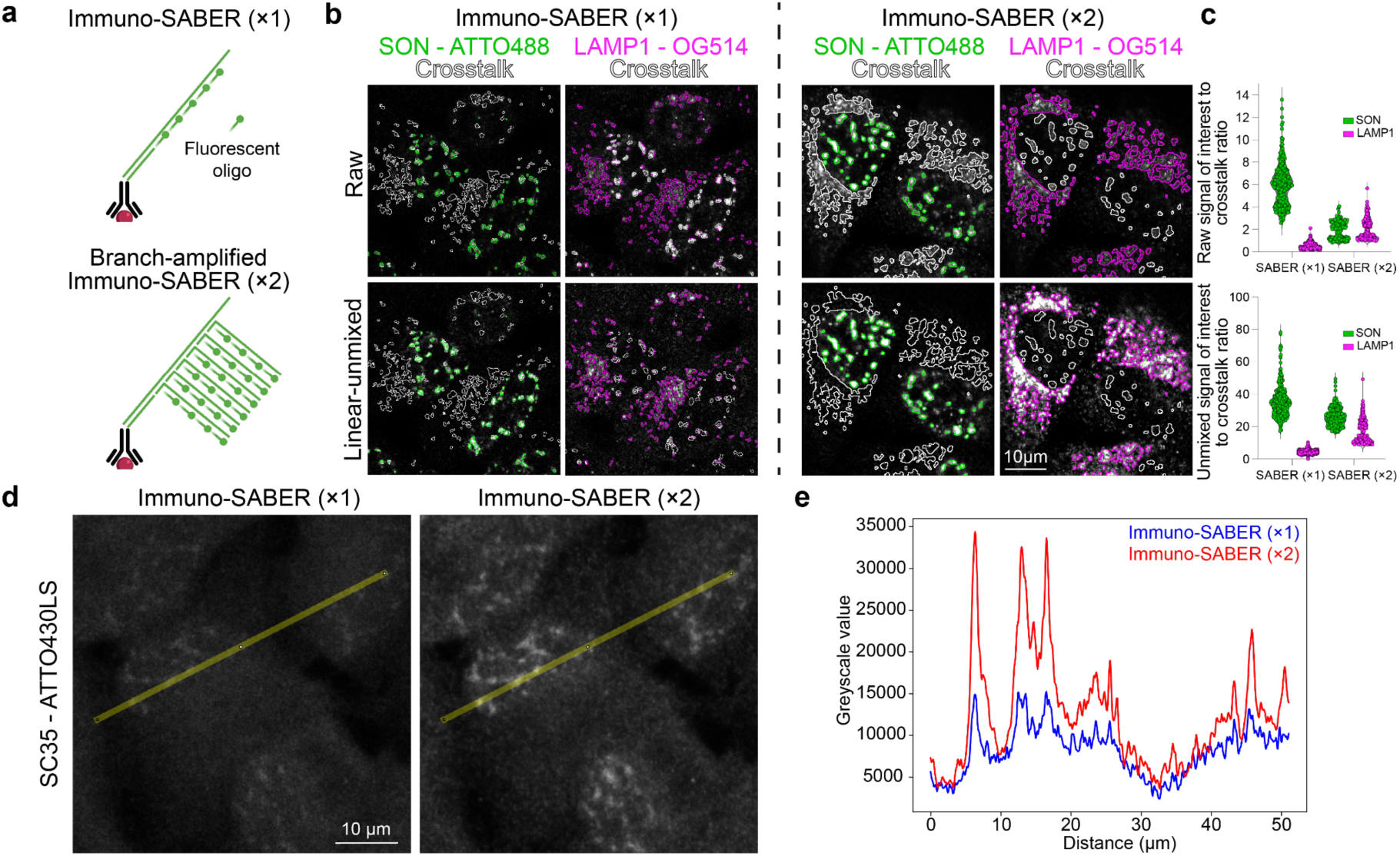
Iterative signal amplification improves overall fluorophore intensities and unmixing of spectrally overlapping dyes. **(a)** Immuno-SABER offers multiple, iterative rounds of concatemer hybridization, thereby increasing the number of potential imager binding sites and target intensity levels. **(b)** Representative images of SON-ATTO488 and LAMP1-OG514 in Immuno-SABER using one round of amplification (×1, linear concatemer) and two rounds of amplification (×2, branching) in the raw and linear unmixed datasets. Green and magenta outlines indicate the signal of interest and white outlines indicate crosstalk in each comparison. **(c)** Upon branched amplification SON-ATTO488 and LAMP1-OG514 signal to crosstalk ratio evened out in the raw dataset, which in turn increased LAMP1-OG514 signal to crosstalk ratio in the unmixed dataset. Note that different imaging settings and unmixing matrices were used for ×1 and ×2 amplification conditions (reoptimized to account for signal differences in both settings), so the signal intensities are not directly comparable (indicated by a dashed line) and are displayed with individual scaling (settings are described in **Supplementary File 2 - Tabs 2-4** for Immuno-SABER ×1 and **Tabs 6** and **7** for Immuno-SABER ×2). n = 175 - 656 ROIs from 2 FOVs. **(d)** HeLa cells stained for SC35-ATTO430LS with Immuno-SABER ×1 or ×2 branched amplification. Acquisition settings are described in **Supplementary File 2 - Tab 12**. **(e)** The final signal intensities obtained with the same imaging settings were compared across the line plot over the same nuclei (yellow line in **d**).

Motivated by this improvement, we also tested whether signal tuning can help us to utilize channels that did not yield high enough signal to be safely unmixed or assign optimal detection parameters. An example of this was SC35-ATTO430LS (a nuclear speckle marker, whose distribution largely overlaps with SON), which under spectral imaging conditions had very low signal (**Fig. 4d**, Immuno-SABER ×1). By performing branched amplification, we doubled the SC35 signal intensity across the same nuclear speckle regions (**Fig. 4d-e**) and recovered the expected staining pattern for this target and hence we were able to incorporate this target-fluorophore combination in our panel as the 15^th^ marker for the following experiments.

### Detecting perturbation-induced changes in subcellular organization

Next, we applied our optimized final panel and imaging conditions in the context of a perturbation experiment. We applied two chemical perturbations (Sodium Arsenite and Actinomycin D) that are expected to affect the visual phenotype of the cells significantly. Sodium Arsenite (NaAsO_2_, SA) is an environmental toxin that exerts various effects on cellular functions, primarily through the induction of oxidative stress. It is well-known for inducing formation of stress granules and decreases overall translation. Hence, we expected a drastic change in the G3BP1 distribution, going from the diffuse cytoplasmic pattern to bright cytoplasmic aggregates. Actinomycin D (ActD) inhibits RNA synthesis by intercalating into DNA^41^. It has a strong effect on RNA Polymerase I and rRNA transcription, so is known to result in loss of nucleoli, and in concentrations >1 μg/ml, it can inhibit all 3 RNA polymerases, arrest cell cycle in G1 phase and can induce apoptosis^42–45^. In our experiments, HeLa cells were treated with 5 µg/ml ActD for 5 h or 0.1 mM SA for 1 h prior to fixation. Since both chemicals are expected to alter the spatial localization and signal intensity of selected markers in the panel, we aimed to leverage these conditions to test whether spectral imaging and unmixing can be performed reliably for our optimized panel. The unmixing matrix was generated by using the images of untreated cells as reference, and imaging settings were kept constant across the conditions, with one exception. Due to condensation, G3BP1 yielded very high intensity under SA treatment; so in order to stay within detector linearity, we had to lower the laser intensity for this channel (**Supplementary File 2 - Tab 6**). This reduction in laser intensity was corrected by multiplication of the signal in the corresponding channel before application of the unmixing matrix (also see **Supplementary Note 1**).

We were able to validate the expected phenotypes in the treated cells, without any visually obvious artefacts coming from unmixing (**Fig. 5a**, **Supplementary Movie**, **Supplementary File 2 - Tab 16**). We then further analyzed the multiplexed images more quantitatively to resolve the phenotypic signatures of these perturbations on different structures.

**Figure 5.**
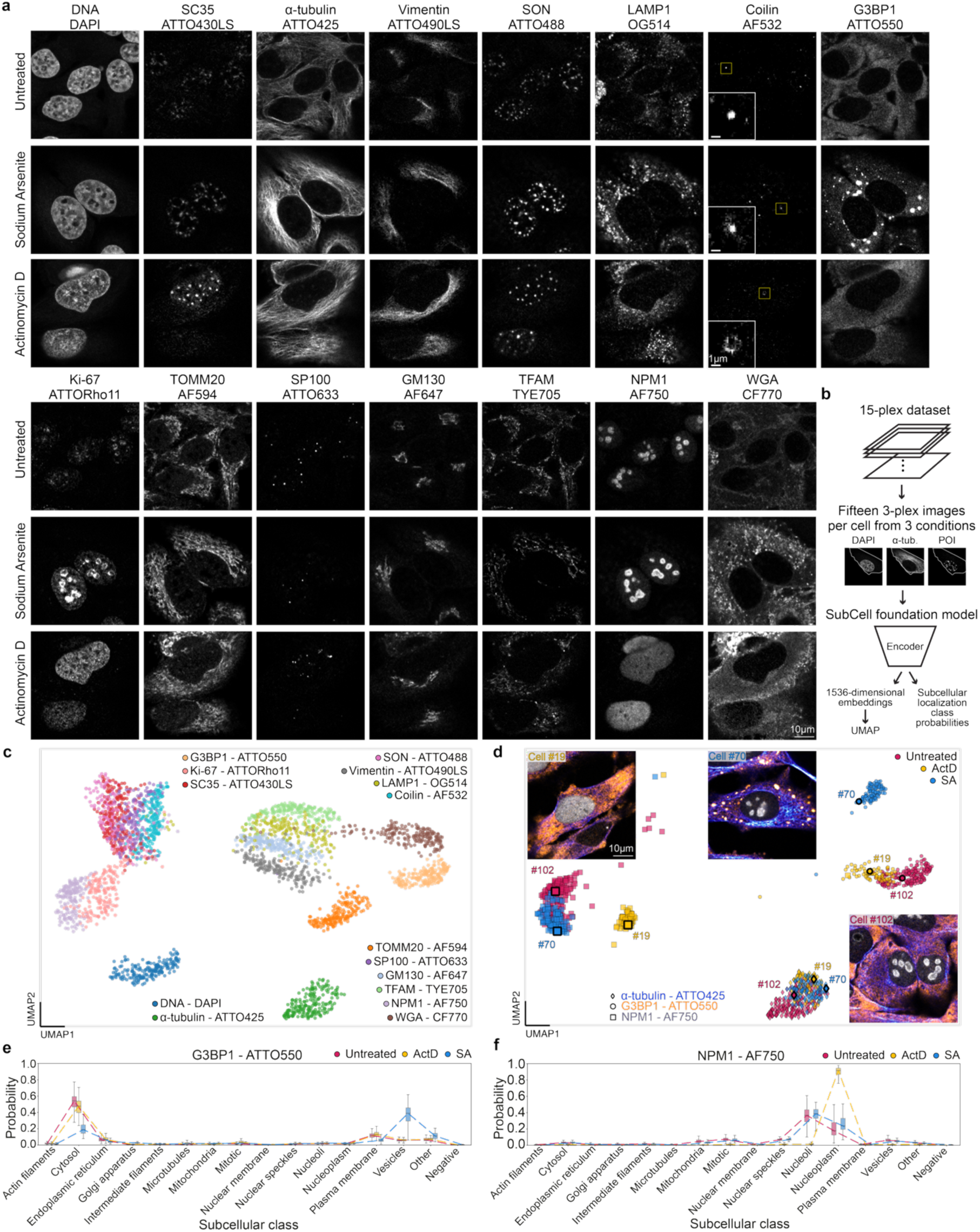
Assessment of the effect of chemical perturbations on the organelle localization. **(a)**Representative single plane images of untreated and perturbed HeLa cells for each protein target after linear unmixing. Untreated cells were separately imaged with three spectrally non-overlapping fluorophore groups as before to generate the unmixing matrix, which was then used for all samples. Insets in the Coilin-AF532 column show a zoom in for Cajal bodies. **(b)** Obtained 15-plex datasets from 3 conditions were split into 3-plex images per cell and subjected to the SubCell foundation model to obtain 1,536-dimensional embeddings and subcellular localization class probabilities. **(c)** UMAP embedding of SubCell output for interphase cells show separate clusters for each organelle for untreated cells. **(d)** UMAP embedding showing the perturbation effect on clusters of selected targets (diamonds: α-tubulin, circles: G3BP1, squares: NPM1). One representative cell is shown for each treatment condition (Untreated: red, ActD: yellow, SA: blue on UMAP) to exemplify the single-cell level link between the clusters and phenotypes for the selected markers. 3-color images show the expected perturbation effect on stress granules (G3BP1, shown in orange in overlay images) and nucleoli (NPM1, gray), without much change for α-tubulin (blue). Shown images are single z-plane images from the 15-plex acquisitions which are processed with linear unmixing. **(e-f)** Evaluation of the relative distribution changes for **(e)** G3BP1 and **(f)** NPM1 by SubCell’s class probability output. Box plots show median probability values per class. Boxes indicate the 25th and 75th percentiles, and whiskers extend to the minimum and maximum values within 1.5 times the interquartile range (outliers excluded). Sample sizes: n=209 (Untreated), 104 (ActD), and 120 (SA) cells across 2 FOVs per condition. Acquisition settings are described in **Supplementary File 2 - Tab 6** and **7**.

Multiplexed organelle-level readouts have proven highly informative for profiling cellular responses to genetic, metabolic, and pharmacological perturbations, capturing coordinated changes in subcellular organization that are not accessible through single-marker analyses. Extending such profiling to higher multiplexing and spatial resolution without relying on multi-cycle imaging would substantially expand the scope of subcellular phenotyping, positioning high-plex spectral unmixing as an enabling strategy for scalable, information-rich subcellular analysis. However, reliable segmentation of each subcellular structure for extracting pre-defined imaging-based features (such as area/volume, number, intensity, morphology, texture), especially under different treatment conditions, remains to be challenging. For highly multiplexed datasets this approach typically requires extensive manual interaction with the data and custom thresholding for each channel and condition, or generation of large datasets and annotated segmentations for training of machine learning models.

An alternative to segmentation-based analysis is provided by segmentation-free subcellular profiling^46^, supported by large subcellular image dataset collections generated by the community over the last years^29,47–49^. Here, a vision foundation model SubCell^50^ was recently introduced to learn and predict protein localization and cellular architecture directly from images. For smaller multiplexed subcellular datasets like ours, SubCell provides the ability to perform segmentation-free profiling out-of-the-box, without additional retraining. However, as SubCell was trained on 3- or 4-plex images from the Human Protein Atlas, its applicability to our unmixed data with higher multiplexing is not a given. We therefore tested whether spectrally unmixed multiplexed images are compatible with the pretrained foundation model and examined whether known biological relationships are preserved in the resulting embeddings.

For this, we first segmented the cells using a vision foundation model of the Segment Anything for Microscopy (μSAM) tool^51^ (see **Methods** for more details) and performed manual corrections and annotation of interphase cells to remove the heterogeneity coming from mitotic cells, and of apoptotic cells with obvious membrane blebs (occasionally observed for perturbed cells).

To match SubCell input, we processed each 15-plex image with linear unmixing into 15 three-channel images, each containing DAPI, α-tubulin, and one protein of interest (**Fig. 5b**). The model generates two outputs from this data for every protein of interest: a 1,536-dimensional embedding and a vector of class probabilities for 30 subcellular structures. To reduce the dimensionality of the 1,536-dimensional embeddings, Uniform Manifold Approximation and Projection (UMAP) was applied. The resulting 2D features were used to visualize whether protein localization patterns match prior biological knowledge and whether perturbations affect them in the expected way. We observed distinct clusters for each structure, where proteins that label similar structures tend to be closer together (e.g., overlapping clusters of nucleolar markers NPM1 and Ki-67, and similarly of other nuclear condensates), suggesting that the embeddings produced by the SubCell on unmixed images preserve protein co-localization patterns (**Fig. 5c**). We then assessed the effect of perturbations, the UMAP showed differential shifts in the embedding space for treatment samples for particular organelle groups, while other markers did not show obvious changes and clustered together across conditions (**Supplementary Fig. S5**). Largest shifts happened for the organelles that are expected to be most affected based on qualitative observations (**Fig. 5a**) and literature, indicating retention of biologically meaningful signals. Taking a closer look at G3BP1 and NPM1 (**Fig. 5d**), we observed that the model was able to distinguish the differential effects of ActD and SA from each other, with SA creating the biggest shift for G3BP1 while ActD particularly affects NPM1. α-tubulin clusters stayed together across conditions, as expected for a target that does not get altered by either of the treatments. To get a more interpretable view of the treatment effects, we used the subcellular localization class probability vectors generated by SubCell as a second output. Several subcellular classes that have similar distributions were merged to make visualization easier (see **Methods**). Relative changes of class probability outputs were consistent with expected subcellular localization trends, suggesting that spectrally unmixed images retain recognizable compartment-specific features, as demonstrated by the shift of the G3BP1 probability mass from class “cytosol” to class “vesicles” under SA treatment reflecting formation of the stress granules (**Fig. 5e**). Similarly, NPM1 probability was shifted from “nucleoli” to “nucleoplasm” only under ActD treatment, consistent with the expected nucleolar disintegration effect of ActD^45^ (**Fig. 5f**).

Together, these results show that spectrally unmixed multiplexed fluorescence images can be used as inputs for pretrained subcellular models to generate segmentation-free representations without model retraining and without requiring new large training datasets. Notably, these models were trained on conventional images acquired without spectral unmixing, indicating that linear unmixing preserves the image features required for effective subcellular representation learning. Across perturbation conditions, panel performance remained robust, and the resulting representations enable detailed analysis of perturbation-induced changes in subcellular architecture while bypassing costly and error-prone segmentation steps. Collectively, this establishes spectral unmixing as a practical route to high-plex imaging that is immediately compatible with existing and emerging foundation-model-based analysis workflows.

## Discussion

Multispectral imaging and spectral unmixing approaches have theoretically been available to researchers for decades, yet they have been largely unexploited for routine multiplexed imaging, largely owing to the complexity of generating and validating such panels that can be utilized for robust in situ data generation. Here, we leverage our DNA-barcoded labeling and signal amplification platform, SABER, to offer a generalizable framework for custom panel development and validation for spectral unmixing. Our approach allows decoupling spectral imaging from fluorescent labeling through DNA barcodes, which creates opportunities for efficient testing, optimization and validation of custom high-plex panels. Using cyclic imaging to generate GT images from the same cells with spectrally non-overlapping panels allows for the surveying of labeling and image acquisition conditions to fine-tune the panel for best performance and minimal crosstalk. We provide more detailed suggestions about constructing custom panels for spectral unmixing and troubleshooting in **Supplementary Note 1**.

Using this pairing between the GT and unmixed images, we were able to perform a quantitative assessment of the unmixing performance and evaluate the suitability of different metrics. Although very commonly used to evaluate unmixing success, SSIM fails to capture the accuracy of unmixing with respect to the pixel-level spatial distribution of the signal, which is the most critical information biologists would expect to obtain from cellular imaging experiments. In contrast, PCC appears as a high-fidelity metric, which provides a straightforward and easily interpretable evaluation of unmixing performance. The residual crosstalk approach we illustrate for the subtraction of biological correlation between markers makes it possible to identify potentially problematic channels independent of panel-specific considerations. We further show that individual tunability of signal amplification not only allows boosting the signal in weak channels (fluorophore-target pairings), but also helps to tune the signal-to-crosstalk ratio and improve the unmixing performance for robust panels. Programmable signal amplification is hence critical for increasing the plexity of spectral multiplexing.

Here, we used Immuno-SABER as an open-source DIY platform for spectral panel optimization due to these unique advantages. Alternative cyclic imaging approaches (such as CycIF^52^, 4i^53^ or others^54^) could theoretically also be implemented for the spectral panel building framework we demonstrated; however, they do not offer the flexibility that DNA barcoding does for re-imaging the same targets. In addition to SABER, other protocols like CODEX^33^ also make use of DNA barcoding of antibodies, but lack the signal amplification aspect, which may be crucial for antibody panels where directly labeled primary antibodies are used with narrow detection windows to minimize crosstalk, and especially for subcellular level detection. Finally, alternative DNA-based amplification methods like HCR^9^ could be implemented in a similar framework; however, some adaptation might be needed for individual signal tunability and robust re-detection. In general, DNA-based probes have been used for a plethora of multiplexed workflows from single molecule imaging to tissue profiling with success. Nevertheless, there may rarely be minor background issues. In our experiments we came across one such problem, where two particular structures found in cells undergoing mitosis, mitotic spindles and cytoplasmic bridges, showed some weak signal for multiple channels in addition to α-tubulin, which we determined was not related to spectral or sequence-level crosstalk but rather to the very high molecular density of these structures potentially trapping oligonucleotides. Since resolving the source of any unexpected background or crosstalk would be especially important for spectral multiplexing, we use this niche case as an example and include more details on how additional measures such as excitation and emission wavelength scans or fluorescence lifetime can occasionally help to investigate any suspected crosstalk (**Supplementary Note 2** and **Supplementary Fig. S6**).

A particular concern with high-throughput use of unmixing is the potential for large variations in unmixing performance under conditions where signal intensity or distribution changes. To check the robustness of the panel, we have applied two stress treatments that induce such changes in part of the target organelles. We were able to detect such changes robustly in unmixed images both visually and through unbiased subcellular profiling, despite using only one set of reference images from untreated cells and a single unmixing matrix for all conditions. We view these results as highly promising for broader use of spectral unmixing. However, it is important to note that every panel is unique and may pose different challenges under changing experimental conditions. In this context, the framework we provide for easy generation of GT and the associated metrics would be valuable for the identification of the most problematic channels in the panel to estimate the risk of crosstalk and optimize the design and acquisition settings accordingly to maximize the robustness. For cases where extreme variations in signal are expected for the highest risk channels across conditions, it might be necessary to prepare condition-specific references for linear unmixing. Alternatively, the use of reference-free unmixing approaches could simplify the acquisition and are expected to be more tolerant to intensity changes across conditions (**Supplementary Note 1**). In this regard, our quantitative assessment of a reference-free semi-blind unmixing approach was very promising, with almost comparable performance to linear unmixing (**Fig. 3c-d**). Accordingly, further refinement of available reference-free algorithms and their reproducible implementation would significantly facilitate increased adoption of spectral multiplexing for high-plex cell and tissue biology applications.

The organellar panel developed to showcase the framework was selected to span diverse subcellular contexts, staining patterns, and levels of spectral overlap, making it broadly applicable for benchmarking across different samples. Beyond validation, this panel enables the analysis of changes in subcellular organization across cell types and treatment conditions, providing potential readouts of cellular states, and perturbation/drug responses^53,55,56^. As the number of probed targets increases, however, segmentation-based analysis becomes increasingly difficult, motivating the exploration of machine- and deep-learning-based analysis strategies, including emerging foundation models. A key open question is whether artifacts introduced by high-plex spectral imaging and unmixing could limit the applicability of such models, given that most of the available training data consist of low-plex images. Here, we provide the first implementation demonstrating that spectrally unmixed multiplexed fluorescence images can be used directly as inputs for pretrained subcellular models without extensive retraining or the generation of large training datasets. While this establishes feasibility, as spectrally unmixed datasets become more widely available, broader evaluation across additional markers, spectral configurations, and biological conditions could be performed to better delineate and address potential limitations of such inputs.

Since multiplexed protein detection is particularly challenging due to the spatial overlap of many protein targets, in this work, we have focused on imaging of proteins with Immuno-SABER. However, SABER can also be used for RNA or DNA-FISH^26^, and implemented in a similar fashion for spectral multiplexing of nucleic acid detection. In contrast to proteins, the typical distribution of RNAs as discrete spots makes them very amenable to combinatorial barcoding schemes (such as MERFISH^57,58^, SeqFISH^59^, StarMap^60^, HybISS^61^ or their commercialized versions). Integrating spectral multiplexing with combinatorial barcoding would help unlock the simultaneous detection of dozens of targets in a single imaging round. Additionally, the incorporation of lifetime separation^62^ or thermalplex strategies^63,64^, as well as combining spectral unmixing with cyclic labeling approaches can boost the number of targets that can be detected with fewer labeling/imaging rounds and support higher throughput spatial omics assays in the future.

In addition to high-throughput experiments, high-plex spectral imaging would be transformative for thick samples, including thick tissues, organs or organisms, 3D models like organoids, and expansion microscopy samples, where fluidic exchange is difficult and cyclic imaging is slow.

## Methods

### Cell culture

HeLa cells (DSMZ, ACC 57) were grown in DMEM medium supplemented with 2 mM glutamine, 10% FBS and 100 units/ml penicillin-streptomycin. 10,000-15,000 cells were seeded per well in 18 Well Glass Bottom slides (Ibidi, 81817) and grown overnight. The cells were fixed by adding an equal volume of 4% PFA (pre-warmed at 37°C) directly into the medium. After 5 minutes, the solution was replaced with fresh 4% PFA, and the sample was incubated for a further 25 minutes.

### Chemical perturbations

For stress assessment, HeLa cells were treated with 5 µg/ml Actinomycin D (ActD) (Sigma, A9415) for 5 h or 0.1 mM Sodium Arsenite (NaAsO_2_, SA) (Sigma, S7400-100G) for 1 h prior to fixation. Untreated cells underwent only a medium replacement.

### Antibody-oligo conjugation

oYo-Link reagents (AT1002-100ss or AT1002-mIgG1-100ss) were purchased from AlphaThera with custom Immuno-SABER barcodes^27^ listed in **Supplementary File 1 - Tab 4**. They were individually reconstituted in nuclease-free water according to manufacturer’s instructions. Off-the-shelf antibodies (listed in **Supplementary File 1 - Tab 1**) and oYo-Link custom reagents were mixed (0.6 µl oYo-Link for 1 µg antibody) and UV cross-linked on ice for 2 h. Conjugated antibodies were evaluated on PAGE gels with Krypton and SYBRGold staining. Conjugated antibodies were stored unpurified either at 4°C or -20°C (as glycerol stocks) in their original buffer from the antibody manufacturer. Ki-67 (CST, 9027S) was obtained in custom formulation and conjugated targeting lysine-residues^27^. Conjugated Ki-67 was stored in 50% glycerol, 1× PBS, 0.05%, Sodium azide, 0.375% BSA, 5 mM EDTA and 0.05% Triton X-100 at -20°C.

### Primer Exchange Reaction *in vitro*

Concatemer extensions were prepared *in vitro* as described previously^26,27^. In total, 100 µl reactions (including primers) were prepared with final concentrations of 1× PBS, 10 mM MgSO_4_, 800 units/ml Bst Polymerase, Large Fragment (New England Biolabs, M0275L), 600 µM each of dATP, dCTP and dTTP, 100 nM of Clean.G^28^, 0.15-2.5 µM hairpin in nuclease-free water. First, the reaction mix was pipetted together, excluding the primers (90 µl total volume), and incubated for 15 min at 37 °C. After Clean.G incubation, 10 µl of 10-20 µM primer oligo(s) was added to obtain a 1-2 µM final primer concentration, and the reaction was incubated for another 2 h at 37 °C, followed by 20 min at 80 °C to heat-inactivate the polymerase. Concatemer length was evaluated on 1-2% agarose gels stained with SYBR Gold. Concentrated and purified concatemers were obtained using MinElute (Qiagen, 28004) or Zymo DNA Clean & Concentrator (Zymogen, D4034) columns. Primer sequences and hairpin concentrations used for each reaction are compiled in **Supplementary File 1 - Tab 5 and 6**. Details of the primer and hairpin design criteria have been described previously^26,28^.

### Immuno-SABER

Cultured cells were stained with antibodies following the Immuno-SABER protocol^26,27^ with the following changes:

- Pre-annealing of barcoded ABs with three short complementary oligonucleotides to generate dsDNA barcode. Refer to **Supplementary File 1 - Tab 2** for the sequences.
- Addition of six blocking oligonucleotides in blocking solution to suppress unspecific binding of ssDNA barcodes. Refer to **Supplementary File 1 - Tab 3** for the sequences.
- Addition of decoy oligonucleotides preventing imager crosstalk for iterative staining. Refer to **Supplementary File 1 - Tab 8** for the sequences.

A full list of other available orthogonal oligo sequences can be found in previously published work^27^. Incubations with primary antibodies, primary concatemers, branches and imagers were performed for 2-3 h. WGA-CF770 concentration was increased to 10 µg/ml in 1× PBS, and incubation was extended to 60 min to improve signal strength compared to the amplified targets in the panel. All samples were imaged in 1× PBS. For GT acquisition and unmixing purposes, 1× PBS was supplemented with 1 mM EDTA, 500 mM NaCl, pH 7.4, 1× Trolox, 1× PCA and 1× PCD as described before^65^. A detailed step-by-step protocol will be made available on protocols.io upon publication.

### Panel design

The multiplex panel was designed such that spectrally adjacent dyes targeted non-overlapping organelle markers, alternating between cytosolic and nuclear localization. For more general recommendations on panel design refer to **Supplementary Note 1**. To freely match PER-extended concatemers with any imager (fluorescent oligonucleotide), converter oligonucleotides were mixed in equimolar ratios with the imagers. The converters contain two 20 nt binding sites: one complementary to the concatemer and one complementary to the imager. Refer to **Supplementary File 1 - Tab 7** for the sequences.

The initial GT and 15-plex multiplex imaging datasets were acquired with single concatemer amplification (×1) for all channels. ×1 amplified SC35-ATTO430LS was not used for unmixing and residual crosstalk analysis (**Fig. 3**) due to very low specific signal. During further panel optimization, both ATTO430LS and OG514 signals were amplified using secondary concatemers (branching, ×2) to enhance photon counts and improve unmixing performance (**Fig. 4**).

DAPI and WGA were added to the panel to label distinct cellular organelles (the nucleus and cellular membranes). Additionally, DAPI and WGA served as reference markers for image alignment and cell segmentation.

### Confocal microscopy

Imaging was performed on a STELLARIS DIVE FALCON (Leica Microsystems). Images were acquired with a diode laser at 405 nm and a white light laser with tunable excitation (440–790 nm range) together with a HC PL APO 63×/1.40 OIL CS2 objective (Leica Microsystems). Images were acquired using the LAS X software equipped with SpectraPlex functionality (Leica Microsystems). If needed for fluorophore characterization, lifetime acquisition, and processing utilized the FALCON and FLIM functionalities (Leica Microsystems). Additional images (for **Supplementary Fig. S1b** and **S6d**) were acquired on a STELLARIS 8 (Leica Microsystems) together with a HC PL APO CS2 40×/1.10 Water objective or HC PL APO 63×/1.40 OIL CS2 objective and LAS X software (Leica Microsystems).

### Image acquisition

For similarity/crosstalk analysis (**Fig. 3**), we compared the GT images generated iteratively using the above-described Immuno-SABER protocol with one-shot multiplex images. Therefore, three regions were imaged four times with different combinations of imagers applied. The first three rounds generated the ground truth images of the regions. Each round was a subset of markers of the one-shot multiplex panel. The combination of fluorophores within a round were selected such that there was minimal crosstalk among them. Selection was guided by the “Staining Group” function within SpectraPlex that selects subsets of a multiplex channel set with minimal crosstalk^66^. Different to reference data each round included DAPI as a fiducial stain for later alignment of the iterative rounds. Between each round another set of imagers was applied and the same region was reimaged. In the final round, all markers were applied together.

Detectors were used in the photon counting mode (‘Power Counting’^12^), scanning used a frequency of 400Hz and was performed, if applicable, in the “Frame by Frame” mode. The image format used was 1024×1024 pixels, with a bit depth of 16-bit selected. In all acquisitions the pinhole was set to 1.5 airy units. Detailed acquisition settings listed in **Supplementary File 2.**

Z-stacks in **Fig. 2-4** were acquired using a voxel size of 0.09 x 0.09 x 1 µm. Line accumulation was set to 3 for all sequences. In **Fig. 5**, tilescans with a voxel size of 0.18 x 0.18 x 1 µm were imaged. Line accumulation was set to 3 for all sequences except sequence 2 where an accumulation factor of 6 was used. Z-stack size ranged between 11 and 12 steps.

### Excitation and emission wavelength scan (Λ-λ-scan)

Excitation and emission spectra of α-tubulin-ATTO425 and SON-ATTO488 (**Supplementary File 2 - Tabs 9-10**) were determined using the excitation and emission wavelength scan by LAS X software (Leica Microsystems). The excitation wavelength range was set to 440-790 nm with a step size of 10 nm using the white light laser as light source. The detection range was set to 450-820 and 450-830 nm, respectively, with step size 10 nm and bandwidth 20 nm using a HyDS detector. The excitation and emission spectrum was exported to the “Dye Database” via the “Excitation / Emission Contour Plot” (Leica Microsystems).

### Spectral unmixing

Multiplex images were unmixed using either reference data (“Group”, here referred to as “linear”) or a semi-blind unmixing approach (“Full”, here referred to as “reference-free”) via the SpectraPlex functionality of LAS X^14^. In short, “Group” is a novel approach introduced by SpectraPlex, in which the unmixing matrix is generated using reference data from a set of partially stained samples. LAS X automatically makes suggestions for image regions to be used as references based on SNR, which can be also manually corrected if they are found to be not representing the target structures well. This method deviates from the classical reference-based approaches, which typically require single-stained reference samples.

In contrast, “Full” calculates the unmixing matrix directly from a fully stained multiplex sample using a built-in SpectraPlex algorithm. In addition to the image, the “Full” method incorporates input spectra along with system and setup information.

Reference data acquisition settings and matrices can be found in **Supplementary File 2 - Tabs 4, 5, 7** and **8**.

### Image pre-processing

For **Fig. 2** extended depth of field (LASX 3D, Leica Microsystems) images were processed for image alignment. All images acquired for **Fig. 3** (14 ground truth images, one spectrally mixed (“raw”) with its corresponding linear and semi-blind unmixed versions) were sum projected using z-binning (LASX 3D, Leica Microsystems) prior to image alignment.

For **Fig. 4**, following alignment, SC35 images were first sum projected and Gaussian blur (sigma = 1) was applied. All steps performed in ImageJ (2.16.0/1.54p).

For **Fig. 5**, individual tiles of the tilescans were aligned manually and cropped in z followed by stitching using the “smooth” settings, followed by uniform cropping of 10 pixels to all image borders. For SA condition lower laser intensity was required for G3BP1-ATTO550 to stay within detector linearity^12^. This reduction was corrected by multiplication of the corresponding channel before application of the unmixing matrix with LASX 3D viewer (Leica Microsystems).

### Image alignment

DAPI stain was used as a landmark for channel alignment using *pyStackReg*^67^, a Python implementation of the *ImageJ* extension *TurboReg*. Specifically, for each field of view, the DAPI channel image of one round was taken as a reference image. The affine transformation between the DAPI channel images of other rounds and the reference DAPI channel image was estimated using *pyStackReg*. The transformation matrix was then applied to all channel images of the respective round using the *scikit-image* warp function. Following alignment, rows and columns containing only zero-valued pixels across all channels were automatically removed. An additional uniform cropping of 10 pixels was subsequently applied to all image borders.

For additional alignment of individual channels (SC35 linear and branched data sets), the ImageJ (version 2.16.0/1.54p) plugin *MultiStackReg* was used. Using rigid settings, the DAPI channels of the iterative cycles were aligned, and the resulting transformation matrix was subsequently applied to the other channels.

### Unmixing accuracy analysis

GT, raw, linear unmixed and reference-free unmixed images from three fields of view were used to evaluate unmixing accuracy (**Fig. 3**). GT dataset was acquired from a linear Immuno-SABER sample that had ATTO430LS included, which showed very low signal after imaging GT dataset, therefore was removed from subsequent analysis. The following steps were performed using a custom written python image analysis pipeline. A gaussian filter (sigma = 1) was applied on each image. Then, Pearson’s correlation coefficient (PCC) was calculated for each image pair in GT vs. GT, raw vs. GT, linear unmixed vs. GT and reference-free unmixed vs. GT datasets. PCC for each channel was normalized to the respective GT DAPI PCC value to account for potential drift during image acquisition. For example, raw vs. GT α-tubulin-ATTO425 PCC value was normalized to raw vs. GT Group 1 DAPI PCC value, as α-tubulin-ATTO425 was in Group 1 in the GT dataset. To evaluate unmixing accuracy, we calculated residual crosstalk which shows how much of the spectral crosstalk signal is in the image compared to the GT dataset. To calculate residual crosstalk only positive values were considered in GT vs. GT, raw vs. GT, linear unmixed vs. GT and reference-free unmixed vs. GT PCC matrices, because in majority of the cases spectral crosstalk only contributes to the increase of the PCC. GT vs. GT PCC matrix was subtracted from raw vs. GT, linear unmixed vs. GT and reference-free unmixed vs. GT PCC matrices and absolute values were calculated. Then, a sum of the differences was obtained, normalized to 14 and scaled to the maximum value for each channel across datasets. Normalization to 14 was performed because in an extreme crosstalk case all 14 channels would show a maximum PCC value of 1 leading to a sum of 14 for each channel. Mean values of three fields of view were plotted as scatter plots and box plots with whiskers indicating minimum and maximum value. Significance was measured using one-way ANOVA with Tukey HSD post-hoc test (*p* < 0.05 = *, *p* < 0.01 = **, *p* < 0.01 = ***).

### Branched amplification analysis

Raw and linear-unmixed images from GT and quantification dataset were used to compare signal to crosstalk ratio using one round (×1) or two rounds (×2) of amplification using ImageJ (2.16.0/1.54p). Briefly, 1 pixel sigma Gaussian blur filter was applied on linear-unmixed images, automated threshold was set using Moments algorithm and binary masks were obtained for both SON-ATTO488 and LAMP1-OG514 images. 1 pixel radius outliers were removed and 0.05 µm^2^ area filter was applied. Mean gray value of each region of interest was divided by the median value of the crosstalk regions in the image. Same regions were measured in the raw and linear-unmixed images. When signal in SON-positive areas was considered as signal of interest, LAMP1-positive areas were considered as crosstalk in the same image and vice versa. Measurements from all ROIs were plotted as violin plots with overlapping scatter plots.

### Image visualization for the figures

For **Fig. 2**, the brightness and contrast were adjusted and images were inverted in ImageJ (2.16.0/1.54p). The composite figure was assembled in Adobe Illustrator (29.8.1).

For **Fig. 3**, representative images were selected from GT, raw, and unmixed datasets for visualization. Inverted grayscale LUT was applied and minimum and maximum brightness intensity values were set manually.

For **Fig. 4**, following pre-processing of the SC35 images, the same brightness and contrast settings were applied (scaled across 16-bit) and pixel intensities were measured across several cells (pixel width = 5). All steps performed in ImageJ (2.16.0/1.54p). Values were transformed into line plots in jupyterhub (3.6.1). The composite figure was assembled in Adobe Illustrator (29.8.1).

For **Fig. 5**, representative single z-plane images were selected from linear unmixed datasets. Brightness values were set manually for every channel and kept the same across treatments.

### Movie visualization/processing

Representative regions from the perturbation data were processed for movie visualization using LASX 3D viewer (Leica Microsystems). An additional channel, ‘DAPI outline’, was generated by duplicating the DAPI channel, applying a median filter (kernel size 11), thresholding the intensity, and applying a second median filter with the same kernel size. If required, “Fill Holes” was applied. The outline was created using Sobel filtering. Brightness and contrast adjustments were applied uniformly across all conditions. Labels were added with ImageJ (ImageJ 2.16.0/1.54p). RGB values for each channel are listed in **Supplementary File 2 - Tab 16.**

## Cell segmentation

### Image preprocessing and normalization

15-plex tilescans (untreated, ActD, and SA) were processed in Python. Each channel was intensity-normalized by clamping pixel values between user-defined percentiles (min_th, max_th). This step reduced the influence of outlier pixels (e.g., very bright spots) and suppressed background noise. The clamped values were then rescaled to 8-bit. This conversion was necessary for downstream deep learning analysis, as the model used (SubCell) was trained on Human Protein Atlas images, which are stored in 8-bit format.

### Grayscale conversion and segmentation

For cell instance segmentation, relevant channels (DAPI, Vimentin, WGA, α-tubulin) were summed into a single grayscale image, normalized, and input to the segmentation model. Segmentation was performed with the μSAM framework^51^, which integrates the Segment Anything Model^68^ with tools for segmentation and tracking in microscopy. Specifically, the *vit_lm* variant of µSAM fine-tuned on light microscopy images for improved performance was used. The model produced 3D cell masks that were further refined through morphological post-processing (erosion, dilation with a disk kernel, and hole filling). Segmented objects below a minimum area threshold or touching image borders were excluded, retaining only complete cell instances. After these processing steps segmentation masks were manually inspected and further corrections were made when necessary (to correct cell borders or overlaps, remove segmentation artifacts, or distinguish mitotic or apoptotic cells).

### Subcell analysis

Downstream analysis was performed using the *SubCell* foundation model^50^, specifically the variant trained with three-channel inputs (DAPI, α-tubulin, and a protein of interest). For each segmented cell, 15 three-channel images were generated, with the two reference channels (DAPI and α-tubulin) and each of the 15 acquired channels as the protein of interest. Because *SubCell* was originally trained on 2D images, our 3D stacks were converted to 2D using sum projection. For every input image, the model produced a 1,536-dimensional embedding representing high-level cellular features, and a 30-dimensional vector corresponding to protein localization class probabilities. UMAP was used for low-dimensional visualization of the first model output using the Python umap-learn implementation with the parameters n_neighbors = 20, metric = cosine, and min_dist = 0.5. For exploratory analysis of UMAP embeddings, the Python library *HoloViews* was used for dynamic inspection of the UMAP embeddings^69,70^.

For the second output, the original inference code was modified to output the 30-dimensional predictions in *softmax* form rather than the default *sigmoid* activation. SubCell model was originally trained on 30 localization classes, and we used the model to create the vectors of localization probabilities to evaluate how the perturbations affect the subcellular distribution of the markers that are known to be most sensitive to these chemicals. To simplify the visualization of the probabilities in **Fig. 5**, either original SubCell class names were used or the subcellular classes were grouped in the following manner: **Mitotic**: “Mitotic chromosome”, “Mitotic spindle”, “Midbody”, “Cytokinetic bridge”; **Nuclear speckles**: “Nuclear speckles”, “Nuclear bodies”; **Nucleoli**: “Nucleoli”, “Nucleoli fibrillar center”, “Nucleoli rim”; **Vesicles**: “Vesicles”, “Endosomes”, “Lipid droplets”, “Peroxisomes”, “Aggresome”, “Cytoplasmic bodies”, “Lysosomes”; **Other**: “Centriolar satellite”, “Centrosome”, “Focal adhesion sites”, “Cell Junctions”.

## Data and code availability

The data and code used for analysis have been deposited on Zenodo and GitHub, respectively, and will be made publicly available upon formal publication. In the interim, access is available upon request to the corresponding author.

## Author Contributions

F.F. and S.K.S. - conceived the project.

R.R., K.J. and T.S. - experimental design, sample preparation and imaging data generation.

K.J. and F.H. - unmixing quantification.

T.S., W.M.V., E.P. and H.L. - image processing.

W.M.V., T.H. and E.P. - SubCell analysis.

A.K., W.F., M.J.R. - contributed to supervision.

S.K.S. - experimental design, data interpretation, manuscript drafting and supervision. All authors contributed to writing of the manuscript.

## Acknowledgements

This work was supported by core funding from European Molecular Biology Laboratory and a research sponsorship from Leica Microsystems, and is facilitated by EMBL Imaging Center. This project has received funding from the European Union’s Horizon 2020 Research and Innovation Program under the Marie Skłodowska-Curie grant agreement No 945405 (ARISE, awarded to KJ) and is supported by the Health + Life Science Alliance Heidelberg Mannheim and received state funds approved by the State Parliament of Baden-Württemberg. EP acknowledges EMBL Corporate Partnership Scientific Program Scientific Visitor Fellowship for financial support. We thank Timo Zimmermann and Jorge Trojanowski for scientific discussions and comments on the manuscript and are grateful to EMBL Advanced Light Microscopy Facility and EMBL Imaging Center for their valuable technical support.

