## Supplementary Information for "Enabling high-plex spectral imaging via DNA-barcoded signal tuning and panel optimization"

**Supplementary Information** includes:

**Supplementary Figure S1.** Introduction of decoy oligonucleotides suppresses potential non-cognate concatemer-imager crosstalk.
**Supplementary Figure S2.** Structural similarity index measure (SSIM) for GT vs. GT, Raw vs. GT, Linear unmixed vs. GT and Reference-free unmixed vs. GT datasets.
**Supplementary Figure S3.** Expected endogenous target overlap in the GT images.
**Supplementary Figure S4.** Matrices used for residual crosstalk quantification.
**Supplementary Figure S5.** Global UMAP visualization of SubCell embeddings from subcellular targets affected by chemical treatments.
**Supplementary Note 1.** Best practices for high-plex panel design and acquisition.
**Supplementary Note 2.** Case study illustrating the resolution of potential crosstalk
**Supplementary Figure S6.** Dissecting microtubule-associated crosstalk in mitotic cells
**Supplementary References**

### List of Supplementary Files

**Supplementary File 1. List of antibodies, oligo sequences (barcodes, primers, hairpins, imagers, blockers)**

Tab 1: Antibodies and structural dyes
Tab 2: Barcode-specific blocking oligonucleotides
Tab 3: General blocking oligonucleotides
Tab 4: Barcodes included in oYo Link custom kits
Tab 5: Hairpins sequences and concentration for primer exchange reaction
Tab 6: Primers for primer exchange reaction
Tab 7: Fluorescent (imagers) and converter oligonucleotides
Tab 8: Decoy oligonucleotides (non-fluorescent imager mimics)

**Supplementary File 2. Additional fluorophores, acquisition settings, unmixing matrices.**

Tab 1: Dyes tested but not included in this study
Tab 2: Ground truth image acquisition settings
Tab 3: Settings 1 image acquisition settings
Tab 4: Settings 1 unmixing matrix settings
Tab 5: Settings 1 unmixing matrix
Tab 6: Settings 2 image acquisition settings
Tab 7: Settings 2 unmixing matrix settings
Tab 8: Settings 2 unmixing matrix
Tab 9: FP Base reference spectra ATTO425
Tab 10: Λ-λ scan α-tubulin-ATTO425
Tab 11: Λ-λ scan SON-ATTO488
Tab 12: SC35 comparison image acquisition settings
Tab 13: Settings Stellaris 8 DIVE image acquisition settings supplementary figure S1
Tab 14: Stellaris 8 image acquisition settings supplementary figure S1
Tab 15: Stellaris 8 image acquisition settings supplementary figure S6
Tab 16: RGB values (LUT) for 15-plex composite movie

**Supplementary Movie. Movie displaying example cells from untreated, ActD and SA conditions** (featuring a crop of the Z-stack from data in main **Fig. 5**).

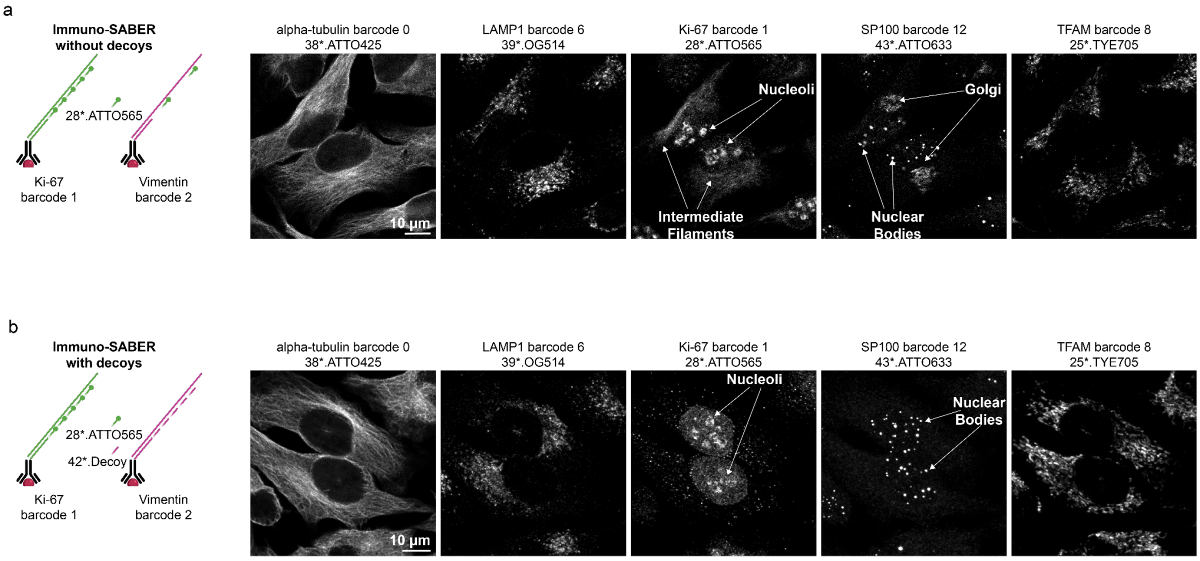


**Supplementary Figure S1. Introduction of decoy oligonucleotides suppresses potential non-cognate concatemer-imager crosstalk**. **(a)** Left: Schematic representation of potential crosstalk, where imagers bind to non-cognate concatemers. In cases where only a fraction of targets from a larger panel are visualized, all single-stranded concatemers serve as potential binding sites. Right: Representative images of five targets (from a 13-plex test panel) simultaneously detected in HeLa cells by their cognate imagers (ATTO425, OG514, ATTO565, ATTO633, TYE705). **(b)** Left: Schematic representation of how decoy oligonucleotides prevent potential crosstalk. Right: Representative images of the group approach using eight decoy oligonucleotides and five fluorescently labeled imagers to ensure specific cognate concatemer-imager pairing. Selected subcellular structures are highlighted with arrows. Settings are described in **Supplementary File 2 - Tab 13** (for **a**) and **Tab 14** (for **b**).

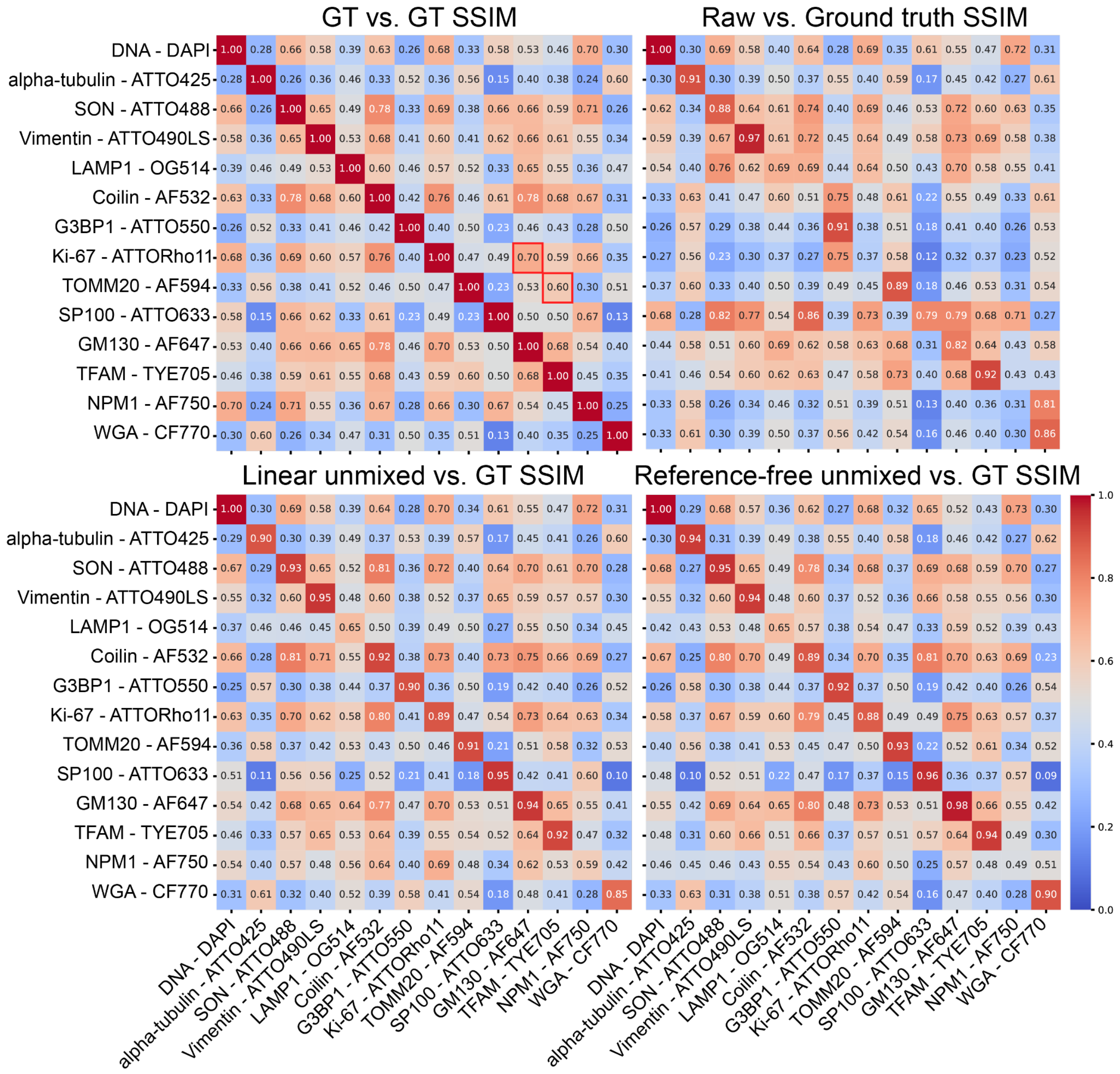

**Supplementary Figure S2**. **Structural similarity index measure (SSIM) for GT vs. GT, raw vs. GT, linear unmixed vs. GT and reference-free unmixed vs. GT datasets**. Red outline indicates unexpected SSIM values in GT vs. GT dataset - high for Ki-67 (nucleoli) and GM130 (Golgi), low for TOMM20 and TFAM (both mitochondrial markers). Numbers on the heatmaps show contain average values from 3 FOVs for each channel comparison.

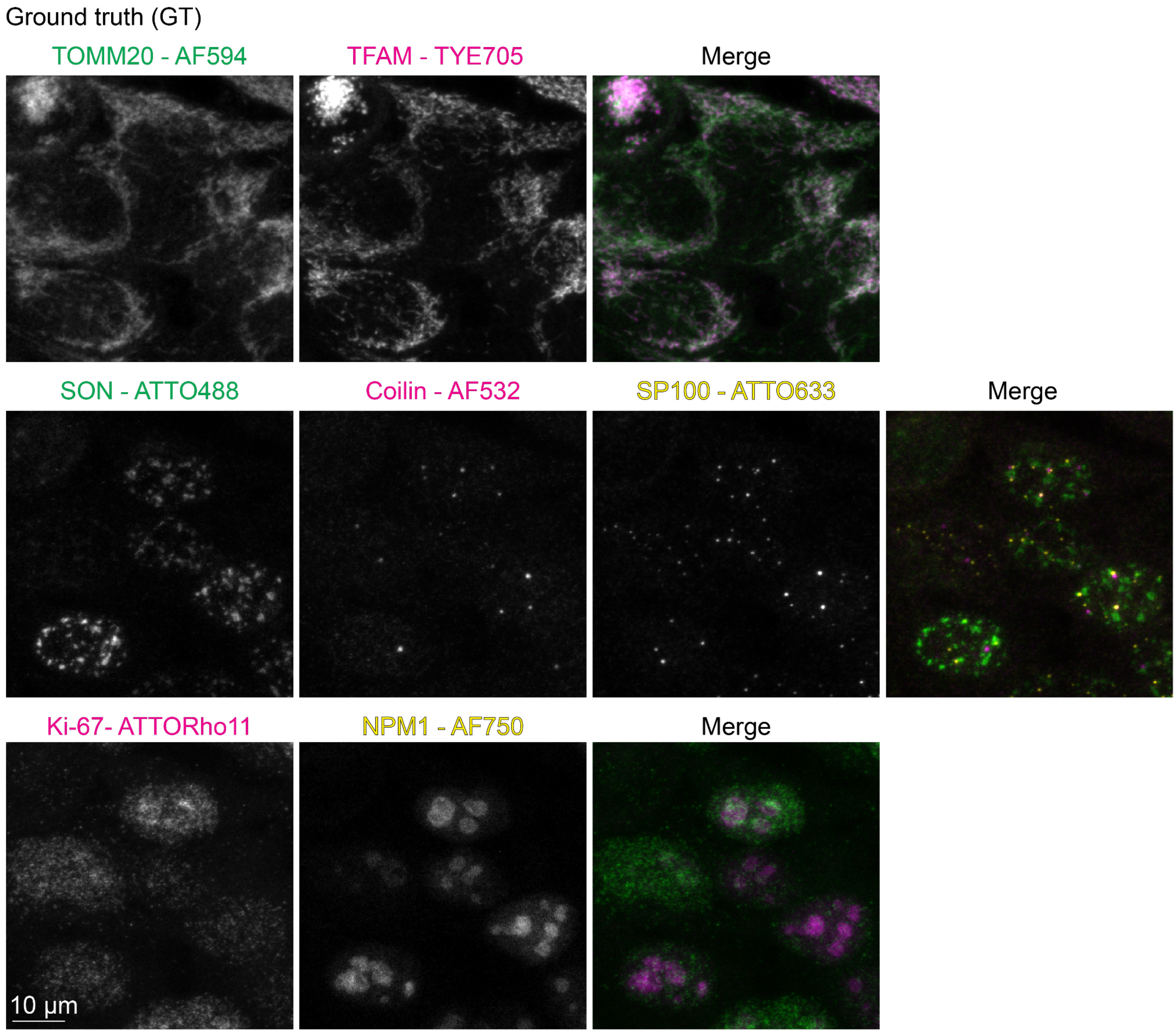


**Supplementary Figure S3. Expected endogenous target overlap for GT images (supplementary to Fig. 3).** Representative sum projection images of targets in mitochondria (TOMM20 and TFAM), nuclear speckles (SON, COILIN), nuclear bodies (SP100) and nucleoli (Ki-67 and NPM1). Acquisition settings are described in **Supplementary File 2 - Tab 2**.

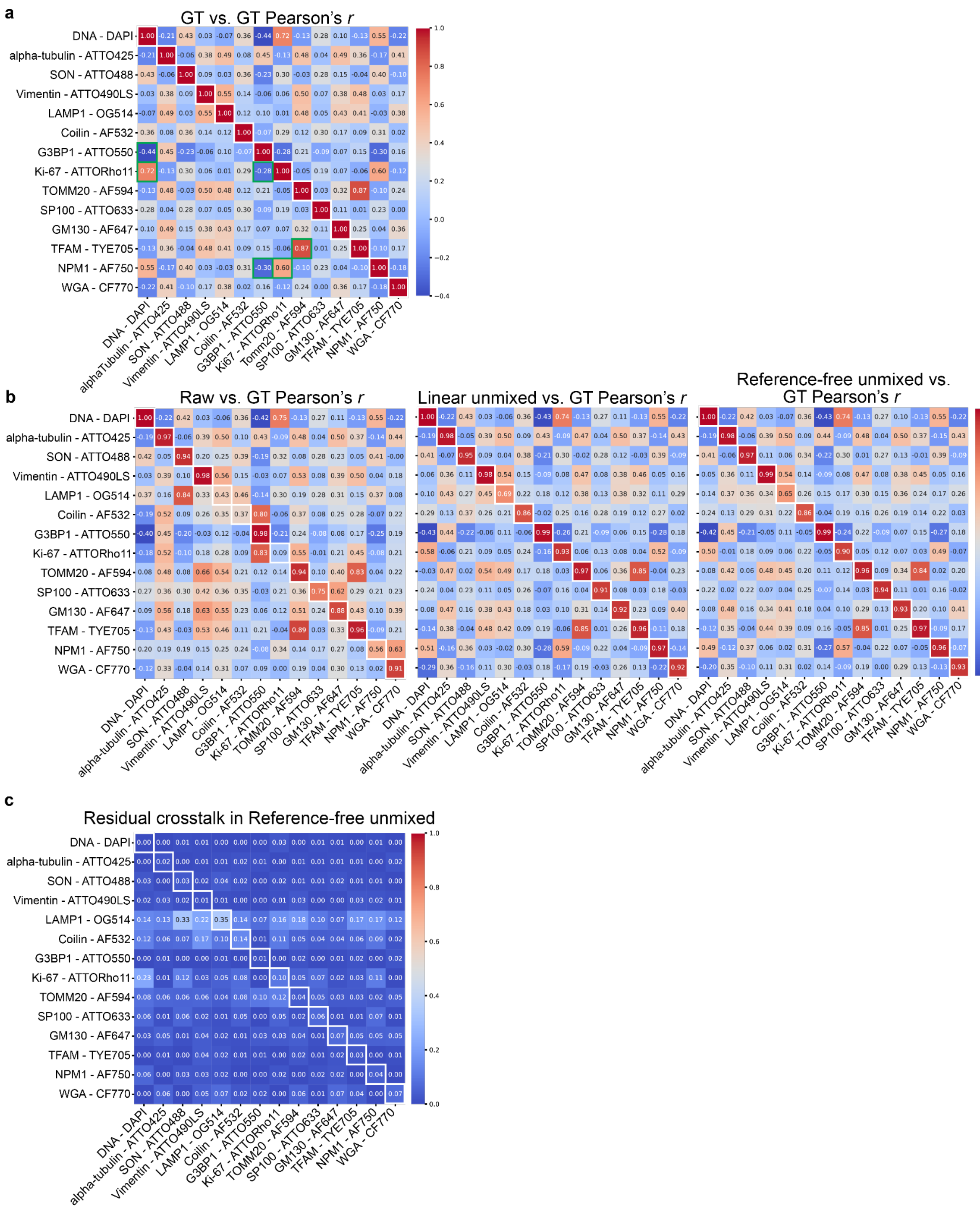


**Supplementary Figure S4**. **Matrices used for residual crosstalk quantification. (a-b).** Normalized Pearson's correlation coefficient for GT vs. GT, raw vs. GT, linear unmixed vs. GT and reference-free unmixed vs. GT dataset (for each channel comparison mean values from 3 FOVs are shown on the heatmaps). Green outlines indicate expected biological correlation in the GT dataset matrix. White outlines indicate self-to-self comparisons, where each channel is evaluated against its corresponding GT. **(c)** Full residual crosstalk matrix after GT subtraction for reference-free unmixed dataset as for raw and linear unmixed dataset in main **Fig. 3b**.

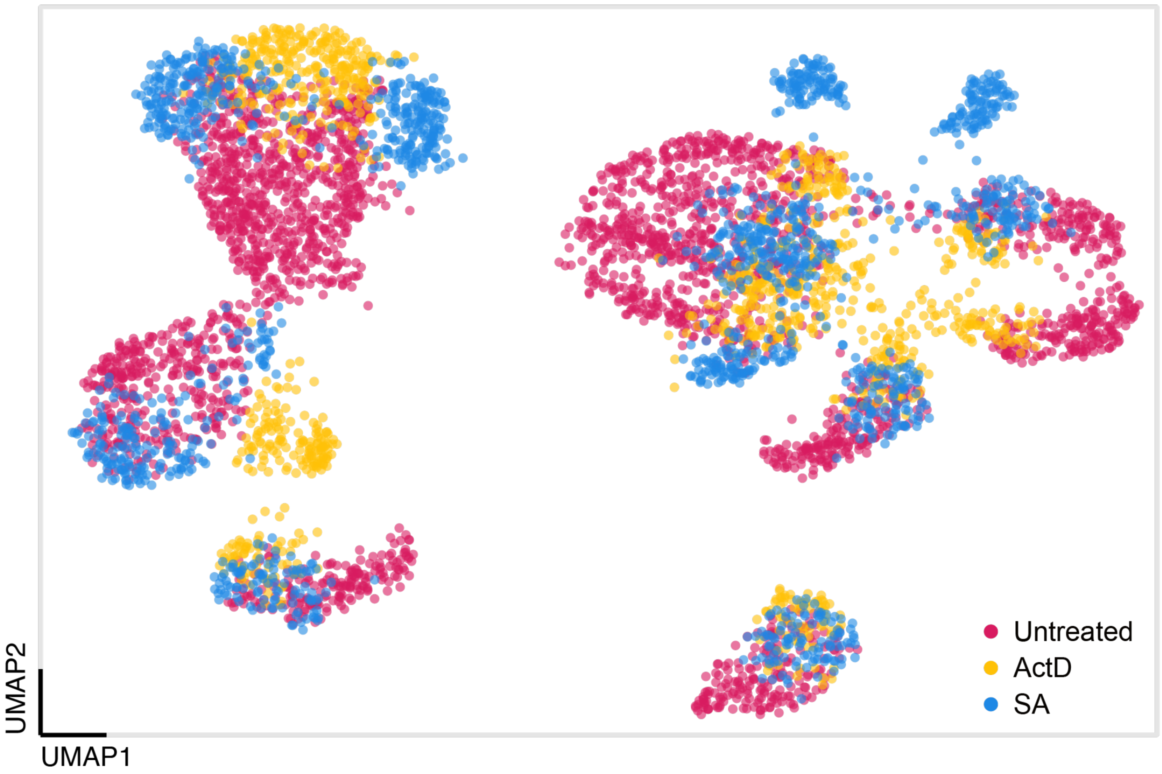


**Supplementary Figure S5. Global UMAP visualization of SubCell embeddings from subcellular targets affected by chemical treatments.** A subset of this UMAP is shown in main **Fig. 5d**.

**Supplementary Note 1. Best practices for high-plex panel design and acquisition.**

| Optimizations/ issues | Recommended actions |
|---|---|
| Fluorophore-target pairing (= marker) | • Generate a hierarchical tree or a co-expression matrix to trace subcellular or cell-type specific expression patterns and as much as possible assign spectrally distant fluorophores to targets that are expected to have highest spatial overlap.<br>• Atlases such as human protein atlas (https://www.proteinatlas.org/)[1] can be used to research target distribution within tissues and cells.<br>• If the expected staining pattern for the target is not known, consider low-plex/standard IF experiments to determine the distribution.<br>• Consider changes in expression to each target under the experimental conditions of interest (perturbations, stress conditions, mutants). |
| Set up the imaging sequence | • Consider simultaneous detection of more than one detector/sequence to reduce imaging time.<br>• Consider combining dyes excited by the same laser line in simultaneous excitation to reduce photobleaching. |
| Set up detectors | • Use detector windows that span over peaks of the emission spectra of the fluorophores in the panel.<br>• Balance the width of the detectors to enable enough photon collection but still be able to see specific signals and not collect too high amounts of crosstalk.<br>• If available, consider photon counting as detector mode for increased sensitivity, brightness, and dynamic range (for example the LAS X STELLARIS offers a Power Counting approach[2], as it simplifies comparison of fluorophore intensities across experiments and reduces noise. Reduced noise helps with more accurate assessment of each species' contribution[3]). |
| Set up excitation wavelength | • Use laser sources that excite at or close to peaks of the excitation spectra of the fluorophores in the panel.<br>• If adequate excitation specificity is maintained, consider exciting at a wavelength away from the excitation maximum. |
| Set up excitation intensity | • Adjust excitation intensities such that all channels show significant specific signal. Direct unmixing with reference-free matrices might help to adjust the simultaneously detected channels such that the expected signal is present in each channel.<br>• If a good balance between excitation intensities of the channels is determined but overall signal intensity is low, consider line / frame accumulation of all detectors.<br>• Avoid saturation from bit depth (change to a higher bit depth) or detector |

| | |
|---|---|
| | linearity (acquire with less excitation intensity, collect less photons by narrowing the detector width, or reduce line / frame accumulation). |
| Acquisition parameters of long Stokes-shift dyes | • Long Stokes-shift dyes (e.g., ATTO490LS) can be added to an already existing high-plex panel due to their unique characteristics of large difference between excitation and emission maximum.<br>• Crosstalk considerations require more close observations since unusual excitation and emission combinations can create crosstalk. This is even more critical since long Stokes-shift dyes tend to have wide excitation and emission spectra.<br> ○ A panel where ATTO490LS is used together with ATTO488 and AF647 can result in high crosstalk of ATTO490LS in the AF647 detector if combined with an excitation within ATTO488 excitation spectrum. |
| Nuclear label DAPI | • DAPI characteristics (405 excitation, wide emission up to 670 nm and high relative brightness) can be incompatible with high-plex panels especially when combined with long Stokes-shift dyes such as Brilliant Violet dyes.<br>• To avoid any DAPI-related crosstalk issues, set up the panel setting accordingly.<br> ○ In this work, potential crosstalk between DAPI, α-tubulin-ATTO425, and SC35-ATTO430LS was avoided by narrowing the detector width to avoid excessive crosstalk and avoid simultaneous detection of these channels.<br>• Alternatively, for image alignment in the optimization/GT part, transmitted light detector (TLD) imaging could be used as the reference channel.<br>• If a nuclear label is required for segmentation and DAPI cannot be used in the panel, consider alternative nuclear labels (such as DRAQ5™, RedDot™ or NucSpot® Nuclear Stains), other segmentation strategies, or segmentation-free approaches. |
| Suboptimal target detection (high background or unspecificity) | • More precise titration of the antibody / detection reagent may be needed.<br>• Alternatively, consult literature, public gene expression and tissue atlas databases to find alternative antibodies for cell type or tissue of choice (e.g., proteinatlas.org)[1,4]. |

| | |
|---|---|
| Cell/Tissue endogenous fluorescence | • If the endogenous fluorescence is defined at a specific wavelength (excitation and emission) it can be handled similarly to an additional dye. In this case, it is recommended to assign an excitation source and detector to capture this signal and image a non-stained sample when generating the unmixing matrix.<br>• If high endogenous fluorescence is expected but wavelength is not known, determine wavelength range prior panel design. Image non-stained tissue with the planned imaging set-up or perform a lambda-lambda scan (Λ-λ-scan), which acquires a full lambda stack (emission spectrum across wavelengths λ at each pixel) while rapidly stepping through multiple discrete excitation wavelengths (Λ).<br>• Skip wavelength range overlapping with endogenous fluorescence.<br>• Detecting additional channels simultaneously with a 405 nm excitation line may lead to endogenous fluorescence appearing in those channels. In this case, evaluate if moving affected detectors to a setting without a 405-excitation line helps to reduce the effect.<br>• Apply bleaching protocol or quenching agent (such as Vector TrueVIEW, Sudan Black B) to reduce endogenous fluorescence.<br>• Assign highly expressed and structurally different targets to the affected channel.<br>• Assign less relevant targets to the affected channel (e.g., channel not required for analysis but for structural orientation).<br>• Amplify target signal with branched SABER.<br>• Endogenous fluorescence may arise from reagents applied e.g., during fixation. This should be checked and avoided if confirmed. |
| Fluorophore performance issues | • Consider alternative fluorophores with similar excitation and emission profiles (for example using FPbase[5,6], vendor catalogues or built-in databases in the instrument software such as dye database of LAS X).<br>• Research extinction coefficient and quantum yield for dyes and assign them to targets accordingly. Pair low expressed targets with high extinction coefficient and/or high quantum yield fluorophores (databases like FPbase display scaled spectra that helps to evaluate these fluorophore properties).<br>• If high crosstalk between fluorophores is expected due to their spectral properties, verify whether their quantum yields and extinction coefficients are compatible and still allow reliable signal discrimination. |
| Fluorophore stability | • Apply oxygen scavenger buffer to sample prior to image acquisition to increase fluorophore stability. |

| | |
|---|---|
| Accuracy of fluorophore spectrum | • Reference-based unmixing matrix generation is rather robust even when the published spectra from the vendor differ from experimentally measured spectra, as long as the spectral alterations remain compatible with the anticipated panel. Minor changes in spectral profile or intensity usually do not affect unmixing, as long as fluorophores remain identifiable and distinguishable. However, acquisition parameters may need adjustments when photons are not collected or excited at the fluorophores' peak wavelengths.<br>• Non-reference-based unmixing matrix generation is considerably more sensitive to discrepancies between published and real spectra, as it relies entirely on the accuracy of the measured data acquired under the specific experimental conditions. If fluorophores are thought to show discrepancy, a Λ-λ-scan should be performed to generate sample-specific excitation-emission profiles. |
| Issues with low intensities of target-dye combinations | • Further amplify target signal with branched SABER.<br>Alternatively:<br>• If available, order fluorescent oligos with multiple modifications of the same dye (eg. on both ends of the imager oligo) to boost the signal further.<br>• Consider swapping to a higher signal yielding fluorophore for this target (see section "Fluorophore performance issues").<br>• Consider performing photon accumulation. |
| Unmixing matrix | • Apply linear unmixing only to images acquired with the same imaging parameters as the matrix was generated. Otherwise, the crosstalk assumptions of the matrix would not be accurate.<br>• Make sure that there were no changes to the instrument setup, calibration or performance (for example instrument maintenance or replacement of a laser) in between unmixing matrix generation and imaging.<br>• If replicates/expected datasets cannot be collected in reasonable timeframe (>6-12 months), consider regenerating the references and obtaining new unmixing matrices. |

| | |
|---|---|
| Unmixing issues in treatments or disease conditions | • Changes of fluorophore intensity are common when treatments or disease conditions alter protein localization or expression level. Even though it is more challenging in high-plex panels, general rules for fluorescent imaging still apply.<br>a. Ensure no channel is saturated (see above) within all considered treatment or disease conditions. This might require determining the range of expected signal intensity of the planned targets.<br>b. If it is not feasible to maintain consistent laser intensities across varying conditions, it will be necessary to regenerate the unmixing matrix by acquiring reference images under the adjusted laser settings. Alternatively, reference-free unmixing methods may be employed.<br>c. As an exception, if detection is linear and the laser does not excite another dye in a simultaneously detected channel, the original unmixing matrix can still be used. In this case, compensate for the change in laser intensity prior to unmixing (i.e., multiply the image intensities with the same laser power scaling factor).<br>• If conditions are expected to create extreme signal intensity or distribution variation, even when the same settings for one panel are used, consider generating condition-specific references and matrices for better unmixing performance. However, the unmixing matrices generated with references where signal in one channel is completely lost, condition-specific matrices cannot be used, and original matrices would be more reliable.<br>• Alternatively, for cases where extreme variation across conditions happen, reference-free unmixing approaches may be considered as they can be more tolerant to variation. However, if the target of interest fails to be yielding the most significant signal in the respective channel (i.e., yield higher signal intensity than crosstalk from the neighboring channels or autofluorescence) then these approaches may result in errors. |

**Supplementary Note 2. Case study illustrating the resolution of potential.**

We occasionally observed unanticipated crosstalk in non-interphase cells for two particular structures: the spindles and midbody (**Supplementary Fig. S6a**). These very microtubule-dense structures unexpectedly showed up in multiple channels (particularly evident in OG514, ATTO633, and AF647), although no spectral crosstalk was predicted based on the excitation and emission spectra of the ATTO425 (**Fig. 1c**), which targets the microtubules. We also performed in situ Λ-λ-scan measurements for ATTO425 and confirmed that the vendor-provided emission and excitation spectra were conserved in our sample *in situ* (**Supplementary Fig. S6b-c**). Lifetime measurements further showed that dyes like AF647 (with reference lifetime of 1 ns[7]) can be detected in these structures in addition to ATTO425 (with reference lifetime of 3.6 ns[8]). Based on these, we confirm and can rule out any spectral crosstalk, hence this was naturally not removed by unmixing algorithms. We did not observe any significant crosstalk in single stainings (for example, GM130 imager does not regularly bind the midbody or spindle structures) or in control experiments, where only one antibody was included but all other components were present **(Supplementary Fig. S6d)**. These controls suggest that the issue does not arise from a direct sequence-level crosstalk between the imagers or concatemers and imagers. Having ruled out these potential sources, we expect that it likely results from unspecific interaction of some DNA oligos with these highly charged and protein-rich structures or a trapping effect that results in ineffective washing of some antibodies or sequences from these very densely labeled meshwork or condensate-like structures. This technical artifact does not affect our results in **Fig. 5**, as our single-cell analysis is focusing on manually annotated interphase cells. For future work, where mitotic cells or these specific structures would be of particular interest, further optimization of the washing protocol to specifically counteract this issue is recommended. We included this example here to illustrate the approach taken here to resolve a suspicious observation to rule out an adverse effect of spectral mixing.

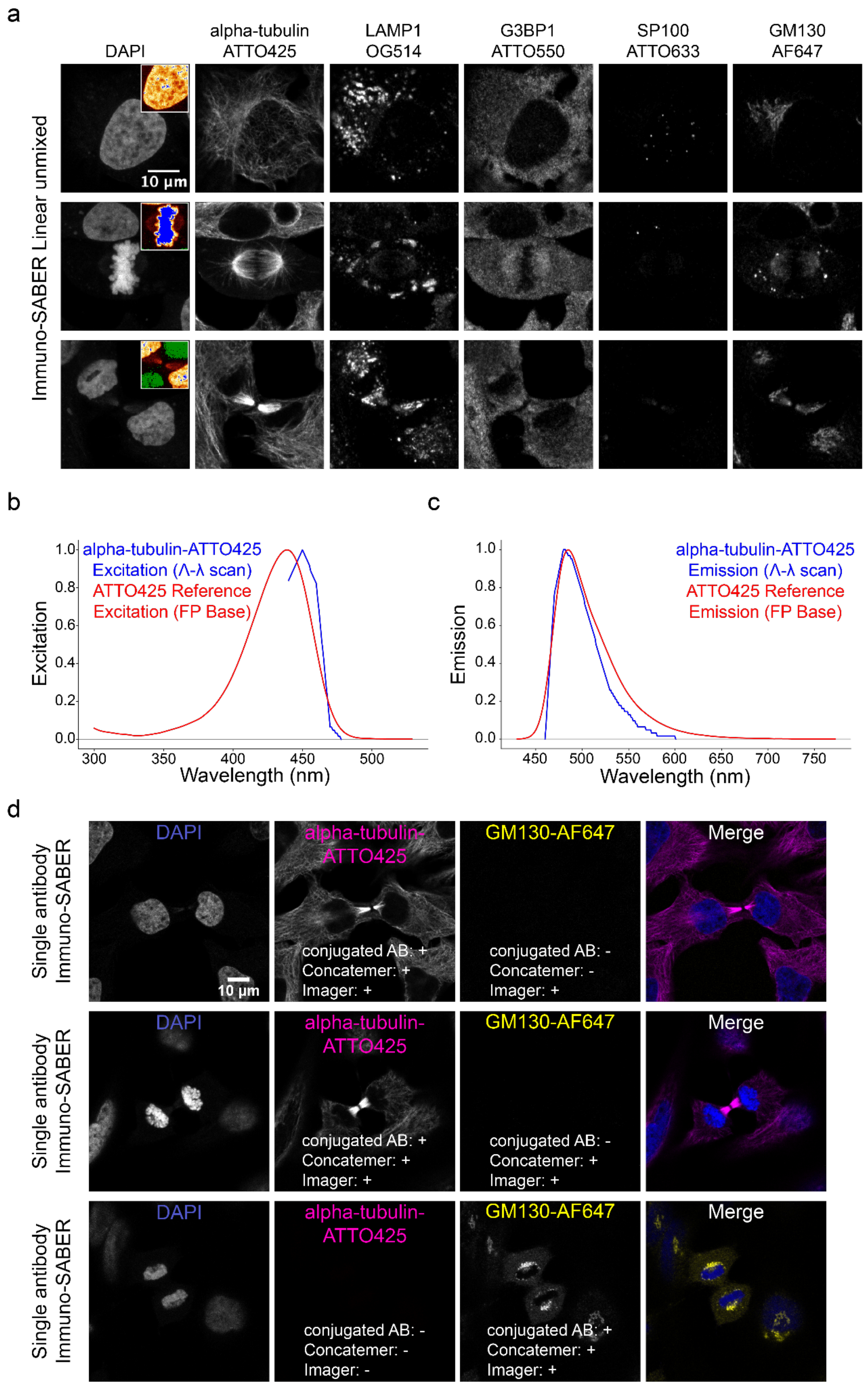


**Supplementary Figure S6. Dissecting microtubule-associated crosstalk in mitotic cells. (a)** Linear unmixing results of the 15-plex subcellular panel in different cell cycle stages. **(b-c)** Comparison of Λ-λ scan (blue) results of samples stained with Immuno-SABER and available reference data[6] (red) for ATTO425 show similar excitation and emission spectra. **(d)** Results of Immuno-SABER for α-tubulin-ATTO425 and GM130-AF647 as single Immuno-SABER stainings in dividing cells. Acquisition settings are described in **Supplementary File 2 - Tabs 6** and **7** (for entire panel **a,** and top and middle row of panel **d**), **Tab 15** (for panel **d**, bottom row). Refer to **Tabs 9** and **10** (for **b** and **c**) for excitation and emission values.
